# Computational simulations of potential *Pachycrocuta* bite damage based on a ∼1.2 Ma ravaged hippopotamus femur from Fuente Nueva 3 (Orce, Granada, Spain)

**DOI:** 10.1101/2024.07.07.602373

**Authors:** Lloyd A. Courtenay, Alexia Serrano-Ramos, Juha Saarinen, Suvi Viranta, Deborah Barsky, Juan-Manuel Jiménez-Arenas, José Yravedra

## Abstract

Understanding the behaviour and interactions of extinct carnivoran species present a significant challenge in archaeological and palaeontological research, often limited by numerous constraints in the fossil record. Here we analyse a hippopotamus femur that presents extensive damage by carnivorans, recovered from the open-air site Fuente Nueva 3 (∼1.2 Ma, Orce, Granada, Spain). Given the extent of furrowing, the potential agents known in the palaeolandscape, and a number of additional theoretical features, we believe this bone to have been consumed by an extinct species of giant hyena; *Pachycrocuta brevirostris*. Leveraging the use of advanced microscopic techniques to digitise the tooth marks observed on this specimen in three dimensions, the present study utilises artificially intelligent algorithms to then simulate the possible morphological variability of this carnivoran. This allows us to propose a characterisation of *Pachycrocuta brevirostris* tooth pit morphology, so as to construct a proposal for a diagnostic reference sample of this species. If our findings are correct, they underscore the importance tooth mark size has on identifying the activity of *Pachycrocuta*, revealing the giant hyena to have produced remarkably large, deep, and circular tooth pits on dense cortical bone.

## Introduction

A complicated challenge in palaeontological and archaeological research is inferring the behaviour of precise agents that have intervened or contributed in the formation of a site. This task is often complicated by our limited ability to distinguish between multiple agents performing either the same, or distinct, activities, yet leaving similar traces. Such intricacies have significantly shaped research surrounding microscopic alterations to bones and their surfaces, fuelling heated debates into the predatory and scavenging capacities of certain species (e.g. Binford, 1981; Blumenschine, 1995; Brain, 1981; Domínguez-Rodrigo et al., 2007, 2021; Espigares et al., 2019; Pante et al., 2012), the dietary habits of others (Domínguez-Rodrigo et al., 2010; McPherron et al., 2010), and the emergence of advanced cognitive tools or abilities within our own evolutionary lineage (D’Errico et al., 2017, 1998; Diedrich, 2015).

Hominins and carnivorans have coexisted and competed for resources throughout the Plio-Pleistocene (Binford, 1981; Domínguez-Rodrigo et al., 2007, 2021; Espigares et al., 2013, 2019; Huguet et al., 2013). The specific dynamics of this coexistence holds significant interest for the scientific community, given that the extent of associated trophic pressures would have profoundly influenced our survival prospects (Turner, 1992). Particularly noteworthy is the southern region of the European continent, presenting a highly complex carnivoran guild during the Epivillafranchian (Antón et al., 2005; Courtenay et al., 2023b; Lozano et al., 2016; Madurell-Malapeira et al., 2020; Madurell-Malapeira et al., 2021; Turner, 1995). Evidence suggests that hominins arrived in this region at least 1.4 million years ago (Arzarello et al., 2007; Carbonell et al., 2008; Huguet et al., 2025; Huguet et al., 2013; Toro-Moyano et al., 2013), thereby providing a compelling landscape for studying these intricate ecological interactions.

One of the keystone species in the carnivoran guild of this region is the giant hyena *Pachycrocuta brevirostris*, documented to have inhabited Europe between *∼*1.9 and *∼*0.8 Ma (Arribas and Palmqvist, 1998; Arzarello et al., 2007; Palmqvist et al., 2011; Madurell-Malapeira et al., 2021; and possibly Werdelin et al., 2023). *P. brevirostris*, along with other hyaenids in general, are posited to have significantly influenced human dispersals across Africa, Europe, and Asia during the Early and Middle Pleistocene (Arribas and Palmqvist, 2002; Madurell-Malapeira et al., 2021). This hypothesis stems from the presumed intense interaction between hyaenids and early *Homo*, supported by numerous sites where both species are evidenced to have been present (Arzarello et al., 2007; Espigares et al., 2013; Huguet et al., 2013; Pineda et al., 2017). Notably, the palaeontological and archaeological sites in the Orce region, situated in the province of Granada, Spain, offer critical insights into this dynamic (Martínez-Navarro et al., 1997; Palmqvist et al., 2011; Toro-Moyano et al., 2013) (Fig. 1). In particular, the *∼*1.2 Ma open-air site of Fuente Nueva 3 (FN3; see Supplementary Materials) has been used to defend the scenario of direct competition between both *P. brevirostris* and early European populations of *Homo* for the consumption of large mammal carrion (Espigares et al., 2013, 2019; Yravedra et al., 2024). Current evidence supporting *Pachycrocuta* activity at this site includes a high prevalence of bones displaying carnivore induced damage in association with abundant coprolites likely attributable to *Pachycrocuta* (Espigares et al., 2023, 2013), and the notably large size of the tooth marks observed on some of the bones. Nevertheless, the Orce palaeolandscape presents a diverse carnivoran guild (Courtenay et al., 2023b; Lozano et al., 2016; Martínez-Navarro and Palmqvist, 1995; Martínez-Navarro 4 Lloyd A. Courtenay et al. et al., 2010, 1997), introducing the possibility that some of these marks may have been made by other significant contributors into the formation of these sites, including machairodontine felids (Martínez-Navarro and Palmqvist, 1995; Palmqvist et al., 2007).

**Figure 1.**
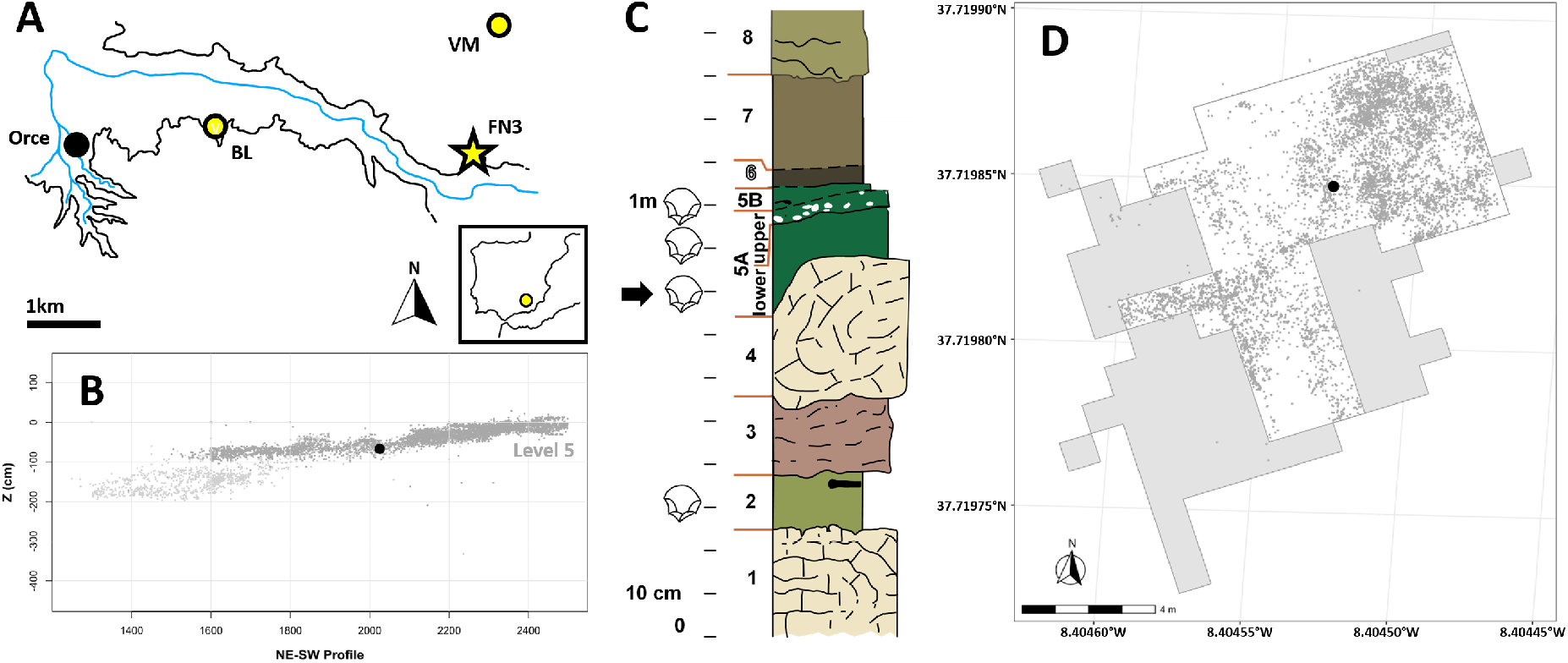
(A) The location of Fuente Nueva 3 in the Orce region, located in the south of the Iberian Peninsula, and its context in relation to other key sites in this region such as Barranco León and Venta Micena. (B) The spatial distribution of fossil finds along the (C) stratigraphic sequence. (D) The geo-referenced spatial distribution of fossil finds recovered from FN3 in the context of the excavated surface. Dark grey areas display the excavated surface in field campaigns prior to 2010 where the coordinates of fossil finds were not recorded. The location of the FN3-11-T93-5-1 specimen is marked by a large black dot in both panels B and D.

While the present authors acknowledge the significant role *Pachycrocuta* would have played in the formation of FN3, pinpointing bones exclusively attributable to *Pachycrocuta* activity, as opposed to other carnivores, remain a challenge (Marean and Ehrhardt, 1995). Recent years have witnessed growing efforts to develop tools capable of distinguishing among carnivore agents based on the bite marks they leave on bones (Courtenay et al., 2021a, 2020). Although some researchers have presented compelling empirical evidence linking specific bite damage to particular carnivoran agents (Courtenay et al., 2023b; Yravedra et al., 2022), such analyses often rely heavily on reference samples composed solely of modern animals, and cannot claim with 100% certainty the agent responsible for producing bite damage. This approach therefore assumes that modern carnivores serve as reliable analogs for their extinct counterparts. Furthermore, unequivocally identifying bones consumed by a particular carnivoran agent remains a rare occurrence (e.g. Domínguez-Rodrigo et al., 2021; Peterson et al., 2021).

Comprehending the behavior of extinct carnivores poses a significant challenge, primarily due to our inability to directly observe them. The focus of this research is to therefore address the inherent ambiguity in identifying *Pachycrocuta* activity amidst a diverse ecological landscape. The primary objective is to establish a robust framework for distinguishing *Pachycrocuta* tooth marks, facilitating the analysis of the trophic pressure within the Orce paleolandscape. Here we present an exceptional example of a fossil likely consumed by a large hyaenid, recovered from the site of FN3. Using sophisticated artificially intelligent algorithms, we use this specimen to simulate the potential variability of tooth pits attributable to the activity of indviduals from the genus *Pachycrocuta*. This research offers a foundational reference that can facilitate the identification of *Pachycrocuta* activity in ambiguous instances of bite damage on bones, while presenting a new perspective for the identification of tooth marks produced by extinct carnivoran taxa.

## Material and methods

### Modern Reference Samples

For the current study, we employed a reference dataset of 823 modern carnivoran tooth pits, sourced from established datasets (Courtenay et al., 2021a,b). Given that the aim of this study is to model variability in tooth pits across different carnivore taxa, it was essential to include a broad and representative sample of carnivore species to ensure that the resulting model is robust and broadly applicable. This dataset encompassed tooth pits generated by a diverse range of species, including bears (*Ursus arctos*, n = 69), spotted hyenas (*Crocuta crocuta*, n = 86), wolves (*Canis lupus*, n = 283), African wild dogs (*Lycaon pictus*, n = 89), foxes (*Vulpes vulpes*, n = 53), jaguars (*Panthera onca*, n = 77), leopards (*Panthera pardus*, n = 84), and lions (*Panthera leo*, n = 82). These tooth pits were derived from both wild and captive specimens, primarily observed on dense cortical bone regions such as diaphyses in medium to large-sized animals. While pooling tooth marks from wild and captive animals can be considered problematic, Courtenay et al. (2021b) demonstrated that for tooth pits this is less of an issue. Digital reconstructions of these tooth marks were initially acquired through high-resolution structured light surface scans (Courtenay et al., 2020). For a comprehensive understanding of the tooth pit characteristics and the digitization protocols, readers are referred to the original publications detailing these samples (Courtenay et al., 2021a,b), as well as the methods (Courtenay et al., 2020).

### Geometric Morphometrics (GMM)

Each tooth pit was characterised using a 25-landmark configuration (Courtenay et al., 2020), comprising of 5 fixed landmarks and 20 sliding surface semi-landmarks. The 5 fixed landmarks mark the maximum length, width and depth of the pit, with LM1 being considered the point along the axis marking the maximum length farthest away from the perpendicular axis marking the maximum width. LM2 is then the opposite point along this axis, LM3 on the left most extremity of the perpendicular axis, and LM4 on the right-most extremity. LM5 then represents the deepest point of the pit. Semilandmarks are then placed using a circular patch over the entirety of the tooth pit, and their final position optimised by minimising bending energy based on the Thin Plate Spline (TPS) approach (Bookstein, 1991, 1989; Gunz and Mitteroecker, 2013). Human-induced margin of error of this landmark model is calculated at 0.14 *±* 0.09 mm (Courtenay et al., 2020). Once landmarks are established, coordinates are superimposed by means of a Generalised Procrustes Analysis (GPA) (Bookstein et al., 1985; Gower, 1975; Rohlf, 1996; Rohlf and Slice, 1990; Slice, 2001). GPA utilises different transformation processes (translation, rotation and scaling) so as to superimpose each landmark, facilitating the quantification of minute differences and variability between configurations. This can either be performed in shape or form, the prior including the scaling process to remove the influence of size, and the latter excluding this process (Rohlf, 1996). Due to prior research revealing the importance of the variable size in the study of tooth pits, superimposition of landmarks were performed in form space (Courtenay et al., 2021a, 2020).

### Unsupervised Computational Learning for Simulating Tooth Mark Variability

For the purpose of simulating and modelling the morphological and morphometric variability that could be produced by a given carnivoran we used a combination of Variational Autoencoder (VAE) (Goodfellow et al., 2016; Kingma and Welling, 2013) and Markov Chain Monte Carlo (MCMC) (Courtenay et al., 2023a; Hastings, 1970; Metropolis et al., 1953) algorithms (Fig. 2). VAEs are unsupervised generative learning algorithms that employ an architecture of neural networks designed to learn efficient representations of complex data by capturing underlying patterns and structures within this data. VAEs consist of two parts; an encoder, designed to compress and embed information into a lower-dimensional feature space, and a decoder, used to reconstruct the original output from this embedded representation. The key component of VAEs are the probabilistic nature of the embedded representation, known as latent space (Kingma and Welling, 2013). VAEs are designed as generative algorithms, with the aim of capturing the diversity of the data so as to produce realistic outputs that are not exact copies of the original data. From this perspective, VAEs learn a probabilistic representation of latent space that represent not only the embedded representation of each individual, but also the possible variability this representation can have. From this perspective, VAEs are powerful tools that offer insights beyond basic dimensionality reduction techniques (Goodfellow et al., 2016).

**Figure 2.**
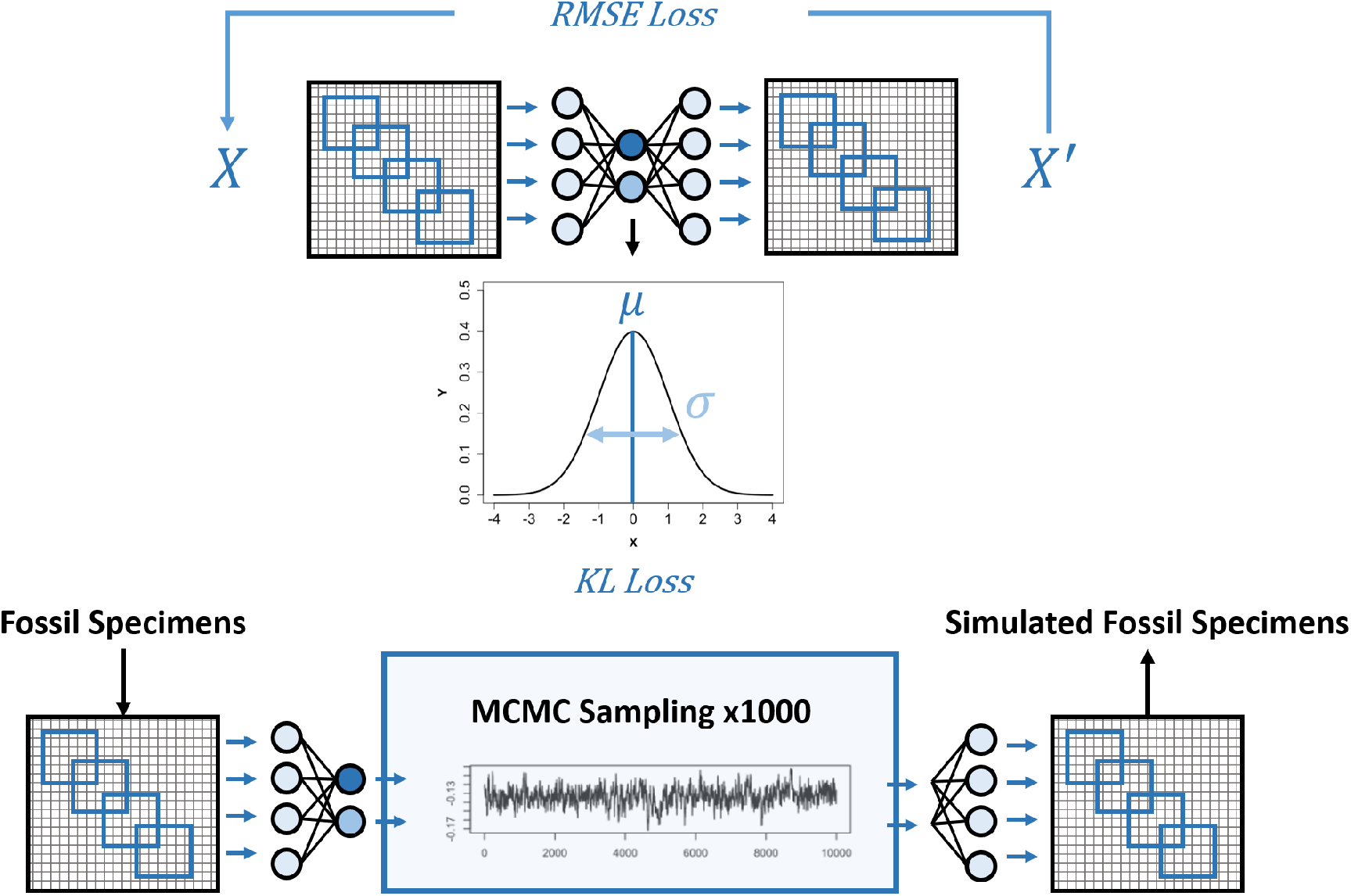
A schematic overview of the algorithm used for simulating tooth-mark morphology in this study. The upper panel illustrates the training of the Variational Autoen-coder (VAE): landmark configurations are passed through a convolutional encoder that reduces them to a lower-dimensional latent space. The decoder then reconstructs the input, and the network’s weights are updated using a combination of reconstruction loss (Root Mean Square Error) and a Kullback–Leibler Divergence term that regularises the latent space towards a Gaussian distribution. The lower panel shows how the trained model is used to generate simulations. The fossil specimen is encoded into the latent space, and a Markov Chain Monte Carlo procedure samples from the resulting probability distribution. These sampled latent vectors are decoded to produce simulated variants of the original fossil tooth mark.

MCMCs, on the other hand, are a family of statistical algorithms that are employed as inference engines in probabilistic programming. MCMCs, as their name suggests, are based on Monte Carlo methods for sampling from probability distributions, yet employ the use of Markovian Chains to stochastically explore a probability distribution (Gamerman and Lopes, 2006). The MCMC therefore learns to sample from a probability distribution randomly by favouring areas of the probability distribution with high information density (Hastings, 1970; Metropolis et al., 1953).

For the purpose of this study, VAEs were trained using 64% (*n* = 526) of the dataset for training, 16% (*n* = 132) for validation, and 20% (*n* = 165) for testing and subsequent evaluation (Bishop, 1995, 2006; Chollet, 2017; Goodfellow et al., 2016). Multiple experiments were performed prior to the final definition of the VAE architecture to test different combinations of hyperparameters and architectures. In continuation we describe the architecture that produced the best results.

VAEs were trained directly on the Procrustes superimposed landmark coordinates using convolutional layers to extract features from them. Both the input and the output to the VAE were thus a 25 *×* 3 matrix of coordinate values. So as to avoid overfitting, the architecture of the VAE was kept as small as possible, consisting in only a single convolutional layer consisting of 64 (3 *×* 3) filters, with a stride of (1 *×* 3), while padding was used to preserve spatial dimensions. For this layer a *L*_2_ kernel regularizer (∥**x**∥_2_) was used (*λ* = 0.0001) (Krogh and Hertz, 1991), as well as a ‘random-normal’ kernel initialiser. Swish was used for the activation function of this convolutional layer, considering swish to perform 0.004mm better in predicting landmarks than other activation functions such as Rectified Linear functions (ReLU: Glorot et al. (2011), He et al. (2015), and Maas et al. (2013)). The output of the convolutional layer was then flattened and passed directly into the latent feature space using a linear activation function. Algorithms were found to perform worse when using both dropout layers and batch normalization (Srivastava, 2013), therefore in the final model these were avoided. The VAE was found to work best when using a latent space of 15 dimensions, serving as a compact and informative representation of the input data. Multiple latent dimensions were considered, however 15 were found to work best. To enable efficient training and gradient propagation, the reparametrization trick was employed (Kingma and Welling, 2013), introducing a deterministic sampling mechanism based on the predicted mean and log-variance of the latent variables. This therefore implies that the flattened output of the convolutional layer is passed into two separate layers of 15 neurons each, one layer to predict *µ* values of the latent representation, and one layer to predict log (*σ*) values of this same representation. This technique facilitates the optimization of VAE parameters, enabling the generation of diverse and meaningful outputs during the decoding process. The latent dimension *z* was therefore defined as (eq. 1);

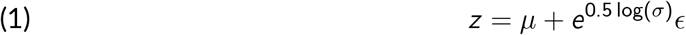

where *ϵ* was additionally found to perform best when sampling from a normal distribution defined as *ϵ ∼* 𝒩 (0, 0.01).

The parameterized latent variables were then fed into the decoder model which was a symmetric representation of the encoder, thus returning the latent variables back to their original structure as a 25 *×* 3 matrix.

The loss function of VAEs comprise of two main components; a reconstruction error and the Kullback-Leibler (KL) divergence loss. The reconstruction loss quantifies the difference between the reconstructed landmarks produced as output and the original landmarks, here using the Square Root of the Mean Squared Error (RMSE). The KL loss, on the other hand, measures the divergence between the distribution of the encoded latent variables and a standard normal distribution (Kullback and Leibler, 1951). This serves as a regularization term that encourages the latent space to conform to a specific prior distribution (previously defined by *ϵ*), promoting the VAE to learn a smooth and structured representation of latent space (Goodfellow et al., 2016; Kingma and Welling, 2013). The KL loss is defined as (eq. 2);

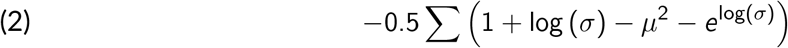

KL serves as the regularization term, encouraging the latent space to follow a standard normal distribution. In the present study, the overall VAE loss was computed as the mean of these two components, balancing the fidelity of reconstructed landmarks with the regularization of latent space during training.

Optimization was performed using the Adam optimization algorithm, and a ‘triangle2’ cyclic learning rate (Smith, 2017), between 0.0001 and 0.01. The VAE was fit for 100 epochs using batch sizes of 32 individuals.

Once the VAE had been trained, evaluations were performed calculating the RMSE between the output of the decoder and the original landmarks provided in the test set. These evaluations found the VAE to have a total reconstruction error of 0.074 mm. Likewise, inspection of learning curves (Supplementary Figure 8) revealed the algorithm to learn well from both the train and validation data with no evidence of overfitting or underfitting.

For the simulation of tooth marks using this algorithm, examples of tooth pits were first encoded using the encoder portion of the VAE, and the corresponding *µ* and log (*σ*) values predicted for each dimension were used to define distributions to be sampled from (Fig. 2). For this, first order variants of MCMCs were used (Courtenay et al., 2023a), utilising the Metropolis-Hastings algorithm as an acceptance criterion (Hastings, 1970; Metropolis et al., 1953). Considering the multiple number of dimensions of the latent space, the probability of a point in multidimensional space was performed by leveraging from the chain rule from probability theory. The MCMC therefore was initialised by taking any random point in latent space. The algorithm then proposed another point in space, and if the probability of the proposed state was considered acceptable, given the Metropolis-Hastings acceptance criterion (eq. 3) (Hastings, 1970; Metropolis et al., 1953), then the MCMC moved to this proposed position in space. If not, then the MCMC stayed where it was.

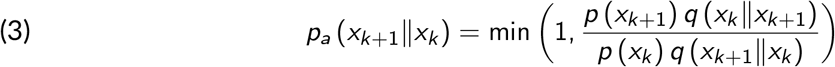

Given the standardised variability observed in latent space thanks to the regularization trick in VAE training, defining a step size for the MCMC was found to be relatively easy. For this purpose, a general step size of 0.005 was set, and adjusted if a particular latent space was found to be more or less difficult to sample from. MCMCs were then used to sample 10,000 points from latent space, discarding the first 2000 iterations for “burn-in”, and then selecting 1000 unique samples from the MCMC’s chain. Evaluation of MCMCs was performed by visually inspecting traces, and calculating Effective Sample (EF) sizes (Kruschke, 2015; Martin, 2018). EF values of above 100 were considered suitable simulations. Supplementary Figures 9 through to 12 present the traces of MCMCs when used to sample from the latent distributions for each of the 4 pits found on the archaeological specimen.

These sampled 15 dimensional points were then passed into the decoder of the VAE to predict 1000 new examples of landmark coordinates based on this compressed representation.

### Archaeological samples and analyses

The tooth marks identified on the FN3-11-T93-5-1 specimen were discovered by one of us (L.A.C.) during the summer of 2021, after a revision of archaeological materials recovered from FN3 during the years 2010 and 2015, located at the Archaeological and Ethnographic Museum of Granada (Spain). Once tooth marks had been identified, silicon moulds of the tooth marks were taken using high resolution silicon (PROVIL NOVO LIGHT), taking the necessary precautions to avoid damaging the cortical surface. Due to the possible accumulative modifications that could be produced to bone surfaces using moulds (Valtierra et al., 2023), a note was left in the fossil’s specimen bag to inform researchers that moulds had been taken if the specimen is to be re-evaluated. The moulds themselves can be inspected upon request to the corresponding author of the current study, and are currently (as of 2024) housed in the University of Bordeaux. Moulds were then taken to the University of Bordeaux for in-depth microscopic analyses.

Tooth pits were analysed directly using the silicon moulds, and then digitised and inspected in detail using confocal microscopy. Specifically, the Mahr MarSurf CM-mobile portable confocal microscope was utilised, housed within the Prèhistoire à l’Actuel: Culture, Environnement et Anthropologie (PACEA UMR 5199) laboratory at the University of Bordeaux (France). This portable device is equipped with a CCD camera with a resolution of 1200 × 1200 pixels, offering a selection of four lenses, providing magnifications ranging from 5 to 100x, all mounted on a motorized platform. This platform features a 50 *×* 50 mm xy-axis and a 35 mm z-axis range, accompanied by a Mahr MarSurf AVDT 120 anti-vibration table to ensure stability during scanning. For this investigation, marks were examined using the Mahr 3200-S lens with 5*×* magnification, characterized by a Field of View (FOV) of 3200 *µ*m, a numerical aperture of 0.15, a working distance of 20 mm, and a maximum measuring angle of 8.6^*°*^. The lateral resolution of the microscope, employing this lens, is 1.93 *µ*m, with a vertical resolution of 1000 nm and a vertical measuring range of 19.9 mm, utilizing white light with a 505 nm wavelength. Scan settings, including illumination intensity (ranging between 90 and 100%), and exposure time (40 milliseconds), were controlled via the MarSurf Metrology SW software, with gain set at 3 db. Image acquisition employed the Standard High Dynamic Range algorithm, while post-processing was conducted externally to maintain standardized parameters across scans.

Three seperate scans were performed to digitise all four tooth pits. These consisted in scans of between 147.6 and 9.46 mm^2^, producing point clouds of between 11.8 million and 20.8 million points, while only an average of 7.7 *×*10^−5^ % points were unsuccesfully measured. Through this we were able to calculate this microscope to succesfully measure 169,957 points per mm^2^.

Following scanning, data files were formatted in FITS and imported into the Mahr MarSurf MfM Premium (v.8.0) software, a part of the MountainsMap software suite developed by Digital Surf (Besançon, France). A systematic workflow was established to create the final 3D models. Initially, point clouds were mirrored to accurately reflect the positive impressions of the tooth pit. Subsequent removal of outliers was achieved through specialized algorithms, targeting isolated artifacts and edge-related artifacts, applied at a moderate intensity. To address non-measured points, a smooth interpolation algorithm utilizing nearest neighbor data was employed, ensuring comprehensive coverage and accuracy in the final model. Point clouds were then imported into CloudCompare, where point normal were computed using a quadric surface approximation model, and an octree radius of 15 *µ*m. The final mesh was produced using a Poisson surface reconstruction algorithm, with a resolution of 5 *µ*m. From meshes, landmark coordinates were then collected and superimposed with the reference samples.

Any additional visual inspection of the cortical surface was performed using a Zeiss EVO 15 Scanning Electron Microscope (SEM) between 50 and 300*×* magnification, utilising the Variable Pressure mode resulting in a chamber pressure of 29 Pa, while using a beam with a 20 kV voltage and 500 pA current.

### Simulating *Pachycrocuta* tooth pits and subsequent analyses

Once both the VAE had been defined and trained on the reference samples, and the landmarks collected from the archaeological specimens, Procrustes coordinates from *Pachycrocuta* were encoded into latent space. MCMCs were then used to sample latent space, and the resulting vectors introduced into the decoder to predict the corresponding landmark values. From the 4 tooth pits recovered from the archaeological sample, 1000 pits were simulated based on each pit, resulting in a total 4000 simulated tooth pits. Marks were then studied metrically and from a morphological perspective using GMM. The metric analyses considered the length, width and depth of the pits, simply calculated by taking the euclidean distances or trigonometric relationships between the fixed landmarks LM1 through to LM5. For GMM, mean-PCAs were computed in form space to compare the *Pachycrocuta* pits with each of the reference samples. This consisted of calculating the central tendency of each species, and using the VAE to predict 1000 variants of these central configurations. Principal Component Analyses were then performed to inspect the distribution of morphological trends, using TPS interpolation (Bookstein, 1989) to predict the morphological changes across each axis of the feature space. Finally, considering PC1 is typically characterised mostly by variations in size, a t-Distributed Stochastic Neighbour Embedding (Hinton and Roweis, 2003) of the PC scores excluding the first PC score was used to assess patterns in the remainder of morphological variables.

### Software

All computational models were implemented in the Python programming language (v.3.10.9), utilising the Tensorflow (v.2.12.0) framework for implementation of VAEs. The numpy (v.1.22.0) library was additionally used for MCMC training. All additional statistical analyses, including geometric morphometric preparation of data, were performed using the R programming language (v.4.3.0), including the geomorph (v.4.0.9) and GraphGMM (v.1.0.0) libraries. All code is available on the corresponding author’s GitHub page at; https://github.com/LACourtenay/VAE_MCMC_Pachycrocuta_Simulations.

## Results

The FN3-11-T93-5-1 specimen (Fig. 3) represents the femoral shaft of a large (Size 5, *sensu* Bunn (1987) and Bunn and Pickering (2010)) artiodactyl mammal. Its overall size and the distinctive cylindrical morphology of the shaft are fully consistent with hippopotamid femora and differ markedly from those of similarly sized perissodactyls, such as rhinocerotids, whose femora typically exhibit a more prominent third trochanter, resulting in a different shaft profile (Bouchud, 1966a,b; Fisher et al., 2010; Pales and Lambert, 1971; Pickford, 2008; Walker, 1985).

**Figure 3.**
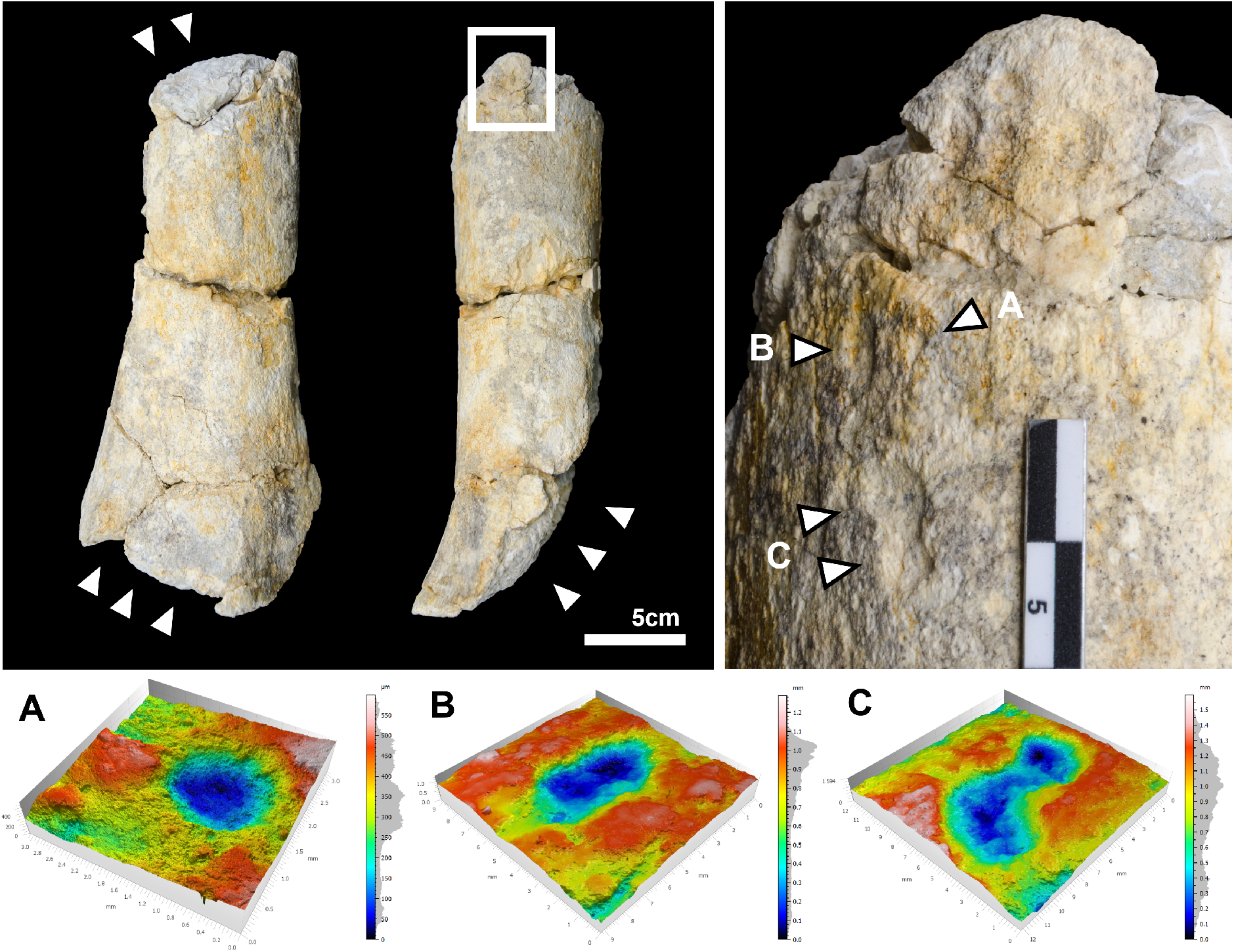
Detail of the FN3-11-T93-5-1 specimen and the identified bite damage. White arrows indicate areas of taphonomic interest. Lower panels present 3D models obtained using confocal microscopy of the 4 tooth pits included within this study (A *n* = 1, B *n* = 1, C *n* = 2). Photos of the Hippo femur were taken by María Higueras Muñoz, figure created by L.A. Courtenay.

To strengthen this attribution, the specimen was directly compared with post-cranial material from extant large mammals housed in the *anatomie comparée* collections of the *Muséum National d’Histoire Naturelle* (MNHN), Paris. Comparative observations were conducted on four individuals of *Hippopotamus amphibius* (two females: MNHN-ZM-AC-1924-134, MNHN-ZM-AC-1943-27; one male: MNHN-ZM-AC-1917-249; and one of unknown sex: MNHN-ZM-AC-1972-308). Additional comparisons were made with femoral remains of several rhinoceros species, including a male *Ceratotherium simum* (MNHN-ZM-2005-297), male and female individuals of *Diceros bicornis* (MNHN-ZM-AC-1944-278, MNHN-ZM-AC-1936-644), and individuals of *Rhinoceros unicornis* (MNHN-ZM-AC-1932-49), and *Rhinoceros sondaicus* (MNHN-ZM-AC-A7971) of unknown sex. None of these rhinocerotid femora exhibit the combination of dimensions and cylindrical shaft morphology observed in the FN3-11-T93-5-1 specimen, supporting its identification as a hippopotamid.

Based on dental remains of hippopotamids recovered from the same FN3 layers (Fig. S1 & S2), this species of hippopotamus for this region and time period is likely to be *Hippopotamus* cf. *antiquus* (Martínez-Navarro et al., 2010), while the dimensions of the femoral shaft also fall within the range of this species (Fig. S3).

Recovered during 2011 field excavations, the specimen was initially assigned the ID number DJ81083 and recorded to have been extracted from Level 4, resulting in its initial label as ID FN3-11-T93-Inf-4-1. In field reports from that year, the specimen was additionally classified preliminarily in the field as a hippo femur. However, subsequent stratigraphic analysis during the 2011 field campaigns revised this association, correctly placing the specimen within Level 5 (Fig. 1b). Likewise, the original labeling system used in previous campaigns would have therefore added the term “Sup” to the label, resulting in the final ID FN3-11-T93-Sup-5-1, however current campaigns do not use the “Superior” and “Inferior” labels for stratigraphy, hence why this part of the ID is omitted here. The specimen in general presents a very poor preservation of cortical surfaces, displaying evidence of biochemical alterations, extensive fluvial abrasion (Fig. S4 & S5), and early stages of weathering. In addition, the medular cavity of the cylinder is filled with a tough concretion. Despite its poor state of cortical preservation, however, this specimen was not subject to intervention by conservation-restoration specialists, and therefore presents limited trephic modifications to cortical surfaces.

FN3-11-T93-5-1 is missing both epiphyses due to intense furrowing of the bone, a feature typical of modern hyena (Brain, 1981; Domínguez-Rodrigo et al., 2015). A number of tooth marks are visible close to the furrowed and crenulated edges of the proximal extremities of this specimen. A total of 7 possible tooth marks have been identified, only 4 of which are distinguishable due to the highly altered state of the cortical surface. Two pits are found in isolation, while another two are found with slight overlap. Nevertheless, the outline of the two pits can still be relatively delimited for the purpose of the present study, and do not mask their overall size.

Despite the challenging state of preservation, all identified tooth pits exhibit considerable dimensions (6.2 *×* 7.7 mm; 4.6 *×* 5.6 mm; 2.0 *×* 2.0 mm; 4.4 *×* 7.8 mm), surpassing the typical range associated with most modern carnivores described by Andrés et al. (2012), and Courtenay et al. (2021a), although does fall into some ranges described by Pobiner (2007).

While we acknowledge that, in principle, tooth marks may be produced prior to, or independently from subsequent furrowing, the quantity of tooth marks observed on the specimen still do not align with typical felid activity (Domínguez-Rodrigo and Pickering, 2012; Gidna et al., 2013)

Considering the improbability of machairodontines causing the observed furrowing (Brain, 1981; Domínguez-Rodrigo et al., 2021, 2015; Marean and Ehrhardt, 1995; Palmqvist et al., 2007), the characteristic durophagous behavior of hyaenids (Palmqvist et al., 2011), the discernible furrowing patterns, and the substantial size of the tooth pits, we can reasonably conclude that the most probable agent responsible for these tooth marks is a hyaenid. Given the current understanding of the palaeontological record of the Orce region, as well as the general understanding of the European carnivoran guild during the Lower Pleistocene, we can be confident in attributing these tooth marks to the genus *Pachycrocuta*.

Utilising VAEs in conjunction with MCMC algorithms, we have been able to model the potential variability of *Pachycrocuta* tooth pits based on those observed on FN3-11-T93-5-1, with a possible 0.074 mm margin of error in predictions. Simulating 4000 tooth pits, our predictions indicate *Pachycrocuta* to be capable of generating notably large tooth pits, with an estimated length, width, and depth, of 6.61 *±* [5.81, 7.37] mm, 4.56 *±* [4.55, 4.57] mm and 0.91 *±* [0.90, 0.93] mm (Table 1), respectively, on dense bone shafts. With the exception of depth parameters, these dimensions exceed the typical range observed in modern hyenas (Fig. 4), and generally surpass those observed in modern lions as well (Andrés et al., 2012). Nevertheless, as with any other modern carnivorans, *Pachycrocuta* had the potential for leaving some relatively small tooth pits as well (Table S1).

**Table 1.** Robust descriptive statistics of the measurements extracted from tooth pit samples, including the simulated measurements of potential *Pachycrocuta* tooth marks. Considering the current species of *Pachycrocuta* identified in the Orce region, as well as the general context of the Western European carnivoran guild during the Lower Pleistocene, we have proposed the attribution of these pits to the species *Pachycrocuta brevirostris*. Measurments are all reported in mm. Upper and Lower bounds of Confidence Intervals (CI) were calculated using 95% interquantile ranges. Detailed description of descriptive metrics with Shapiro-Wilks *p*-values have been included as Supplementary Table 1.

|  |  | Min. | Lower CI | Central | Deviation | Upper CI | Max. |
| --- | --- | --- | --- | --- | --- | --- | --- |
| *V. vulpes* | Length | 0.520 | 1.730 | 1.976 | 0.892 | 2.222 | 3.941 |
|  | Width | 0.501 | 1.12 | 1.41 | 0.593 | 1.520 | 3.209 |
|  | Depth | 0.045 | 0.215 | 0.255 | 0.160 | 0.342 | 1.117 |
| *P. pardus* | Length | 1.272 | 1.977 | 2.410 | 0.972 | 2.744 | 6.346 |
|  | Width | 0.755 | 1.332 | 1.556 | 0.713 | 1.808 | 3.735 |
|  | Depth | 0.038 | 0.222 | 0.266 | 0.207 | 0.347 | 1.493 |
| *C. lupus* | Length | 1.09 | 2.227 | 2.410 | 0.986 | 2.591 | 8.294 |
|  | Width | 0.732 | 1.635 | 1.739 | 0.727 | 1.995 | 4.639 |
|  | Depth | 0.045 | 0.152 | 0.185 | 0.108 | 0.209 | 1.435 |
| *U. arctos* | Length | 0.877 | 2.502 | 3.035 | 1.890 | 3.750 | 9.493 |
|  | Width | 0.496 | 1.584 | 1.975 | 1.172 | 2.562 | 6.695 |
|  | Depth | 0.023 | 0.191 | 0.272 | 0.307 | 0.430 | 2.422 |
| *L. pictus* | Length | 1.125 | 3.231 | 3.482 | 1.703 | 4.480 | 12.596 |
|  | Width | 0.886 | 2.037 | 2.548 | 1.378 | 2.869 | 7.776 |
|  | Depth | 0.059 | 0.384 | 0.497 | 0.442 | 0.612 | 4.160 |
| *C. crocuta* | Length | 1.278 | 3.513 | 3.893 | 1.703 | 4.480 | 12.596 |
|  | Width | 1.096 | 2.495 | 2.776 | 1.290 | 3.097 | 9.444 |
|  | Depth | 0.082 | 0.351 | 0.421 | 0.266 | 0.491 | 3.176 |
| *P. onca* | Length | 0.855 | 3.299 | 3.863 | 2.537 | 4.262 | 16.863 |
|  | Width | 0.621 | 2.093 | 2.437 | 1.713 | 3.092 | 9.947 |
|  | Depth | 0.047 | 0.300 | 0.355 | 0.410 | 0.555 | 2.166 |
| *P. leo* | Length | 2.112 | 4.946 | 5.916 | 3.363 | 6.768 | 18.820 |
|  | Width | 1.241 | 3.688 | 3.994 | 2.209 | 4.818 | 9.680 |
|  | Depth | 0.142 | 0.721 | 0.975 | 0.839 | 1.162 | 3.882 |
| *P. brevirostris* | Length | 1.627 | 5.817 | 6.612 | 2.412 | 7.365 | 8.129 |
|  | Width | 1.589 | 4.546 | 4.556 | 1.613 | 4.566 | 6.533 |
|  | Depth | 0.236 | 0.886 | 0.910 | 0.274 | 0.925 | 1.162 |

**Figure 4.**
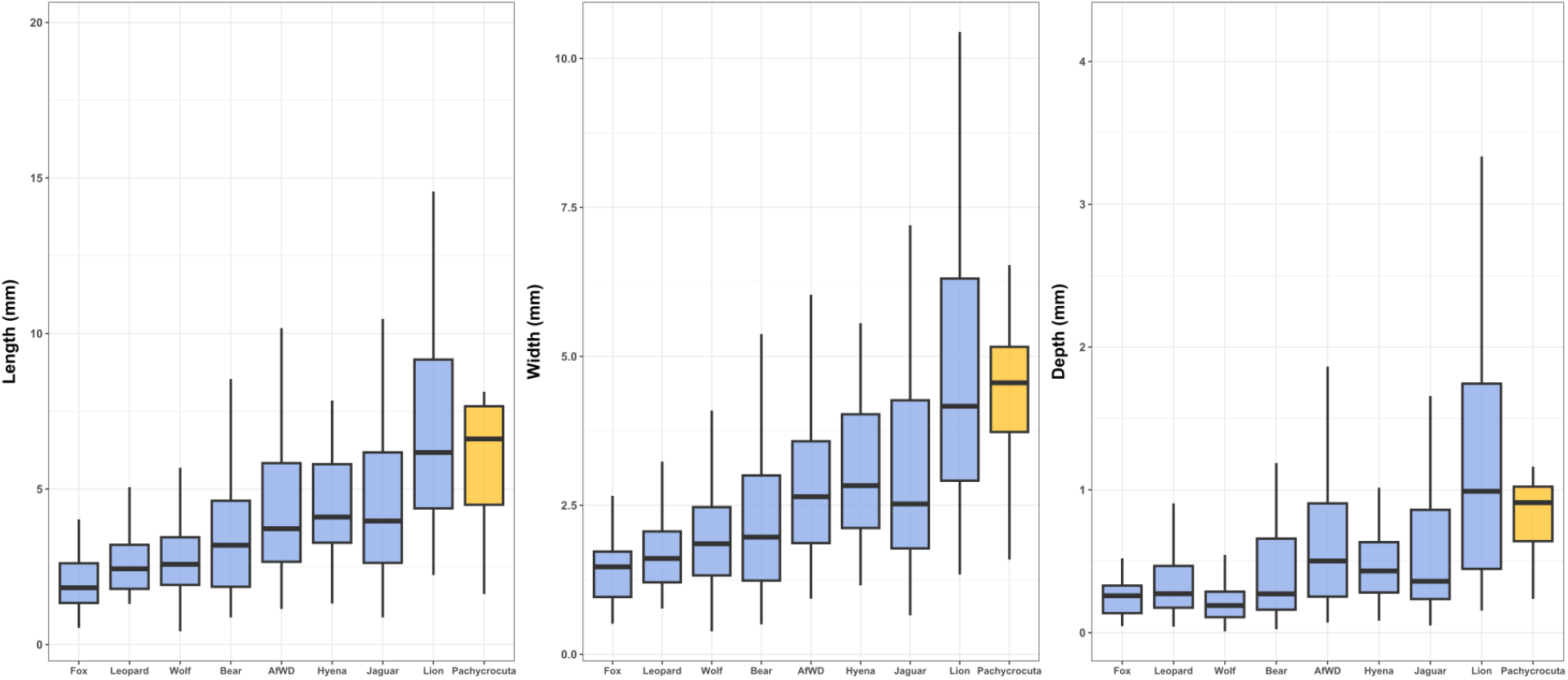
Boxplots visualising the distribution of length, width and depth values in mm of tooth pits produced by modern day carnivorans, as well as the simulated variability that *Pachycrocuta* could have produced. AfWD = African Wild Dog (*Lycaon pictus*).

Upon exploring the possible morphological variability beyond simple measurements of length, width and depth, our proposed sample of *Pachycrocuta* tooth marks have been revealed to produce relatively circular and deep tooth pits (Fig. 5). Combined with their size, these tooth pits therefore appear largely different from almost any modern species of carnivore, yet slightly similar to tooth marks produced by modern lions (*Panthera leo*). These morphological affinities are mostly explained by variation in tooth mark size, which predominates the first dimension of any PCA in form space (Fig. 5), and represents up to 98.31% of the morphological variability here. Nevertheless, our *Pachycrocuta* sample separates from *P. leo* across PCA’s second dimension, which is explained more by the length:width ratio for each pit. In this sense, *Pachycrocuta* tooth marks are more likely to be explained by circular morphologies, where *P. leo* produce more elongated marks. When accounting for the influence of size on tooth pit morphology and focusing on other morphological variations, the proposed *Pachycrocuta* tooth marks remain distinct from the majority of those made by modern carnivores (Fig. S6), although they share similarities in shape with smaller carnivores like foxes and wolves due to the more circular nature of the pits. Across all analyses, another important feature in our simulations is the tooth mark depth, which separates these carnivores from other species producing slightly more superficial pits such as modern felids and canids, and notably hyaenids as well. Nevertheless, the most defining characteristic of this species undoubtedly lies in the size of its tooth pits.

**Figure 5.**
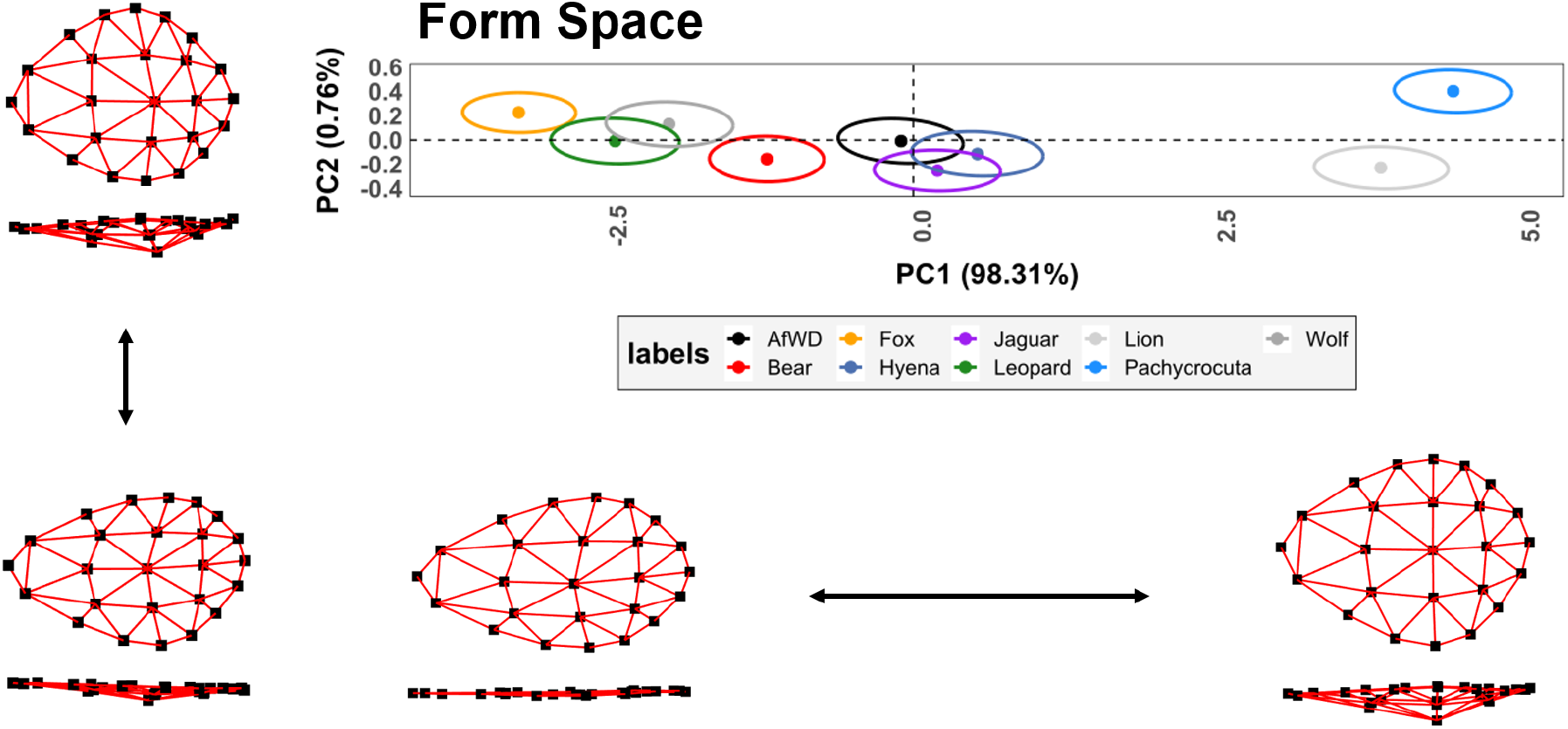
Exploration of the morphological variability among different carnivoran tooth pits in form space using Geometric Morphometrics. Dots indicate the position of the median individual for each species followed by 95% confidence intervals constructed around the median individual through simulations in Variational Autoencoder latent space. Morphological variability has been visualised across the extremities of each axis. AfWD = African Wild Dog (*Lycaon pictus*). Note that our use of the label *Pachycrocuta* refers to the potential marks that could have been produced by this species, as argued in the main body of text.

Finally, to ensure that the results presented are not a product of underlying structure in the VAE latent space caused by sample imbalance, we performed a detailed analysis showing that sample imbalance has no detectable correlation with the structure of the latent space (Fig. 6). A full description of the calculations is provided in Supplementary Appendix B.

**Figure 6.**
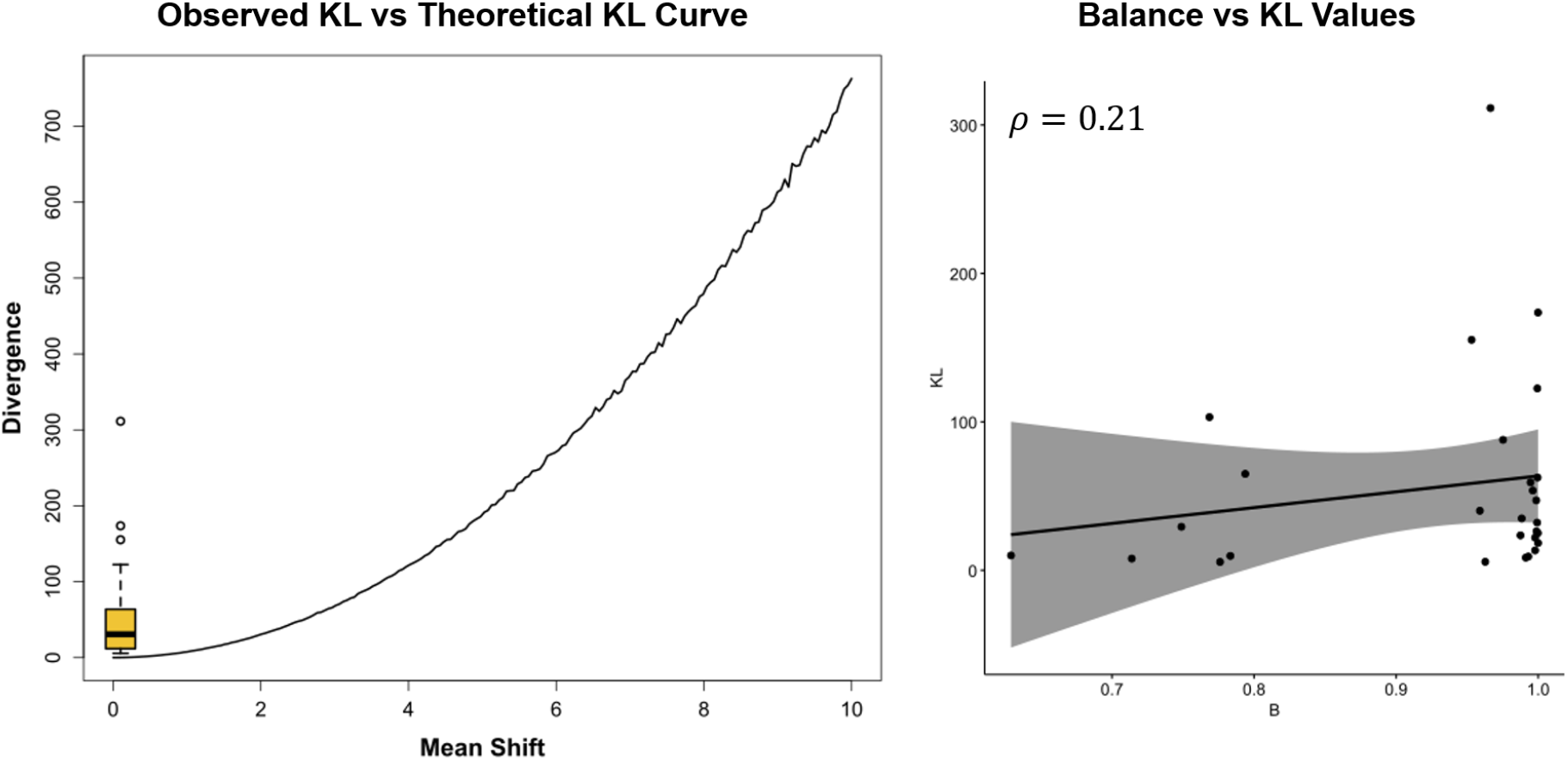
Results of simulations conducted to assess whether sample imbalance and overrepresentation of certain species influence the underlying distribution of VAE latent space. Left panel: Kullback-Leibler calibration curve showing comparisons of distributions with gradually increasing mean-shift values in a 15-dimensional latent space (ℝ^15^). Right panel: Spearman correlation between pairwise Kullback-Leibler divergence values and sample imbalance across species pairs. A full description of the calculations presented here can be found in Supplementary Appendix B.

## Discussion and Conclusions

Identifying the specific carnivore responsible for tooth marks on bone presents a challenge, often influenced by our understanding of modern species, which may not accurately reflect their ancestral counterparts. The presence of *Pachycrocuta*, and more specifically *P. brevirostris*, at the FN3 site is undeniable, evidenced by the abundance of likely *Pachycrocuta* coprolites (Espigares et al., 2023, 2013) and the recovery of numerous dental remains since excavations commenced in the 1990s (Martínez-Navarro et al., 2010; Palmqvist et al., 2011) (Fig. S7). In this context, here we present an impelling example of a bone consumed by hyaenids, confirming *Pachycrocuta* to be the most probable biting agent, given the substantial size of the tooth marks on the bone, as well as the pronounced consumption of epiphyses resulting in a cylinder (Brain, 1981; Domínguez-Rodrigo et al., 2015). Notably, the affected specimen originates from a hippopotamus, a sizable mammal unlikely to be consumed in such a manner by other species. While it is possible for some carnivores, including lions and the modern day spotted hyena, to produce very large tooth pits (Pobiner, 2007), the more fragile dental structure of machairodontines recovered from Orce diminishes their likelihood as durophagous predators (Palmqvist et al., 2007), while other significant carnivore species in this paleolandscape, such as *Canis mosbachensis*, are too diminutive to inflict such damage (Palmqvist et al., 2008). Likewise, *Ursus etruscus* are not particularly known for being consumers of bone nutrients, following a much more omnivorous diet (Medin et al., 2017).

It is important to consider the possibility that remains modified by one carnivoran species could subsequently be scavenged by other carnivorans, potentially resulting in taphonomic palimpsests where multiple agents leave evidence of interaction on the same bone. In Courtenay et al. (2023b) we hypothesised this scenario on a handful of specimens, yet are unable to state with certainty whether this is an accurate identification of such practices. However, in the current case, it is highly unlikely that another carnivoran intervened after the potential activity of *Pachycrocuta*. The extreme degree of furrowing observed strongly suggests that little to no material remained for subsequent modification, and the size of the tooth marks do not fit with any other durophagous agent that could have been searching for nutrients beyond what would have been left after the activity of a hyena. Given the density and morphology of the pits preserved, there is no indication that they pre-date the furrowing. Instead, their distribution and size are entirely consistent with marks formed during the same intensive feeding episode that produced the extensive removal of the epiphysis. A single, continuous consumption event by a durophagous hyena therefore remains the most parsimonious interpretation.

Utilising this exceptional specimen, we have employed advanced artificially intelligent algorithms to produce simulations of potential tooth marks attributable to *Pachycrocuta*. Our models demonstrated remarkable accuracy, highlighting the potential of using both VAEs and MCMCs for the simulation of this type of data. Using this approach we have been able to simulate up to 1000 potential variations for each of the observed tooth pits. It is crucial to note, however, that carnivorans exhibit a broad spectrum of variability (Courtenay et al., 2021a, 2023b), thus not all *Pachycrocuta* tooth marks may manifest to be as circular as those observed on this specimen. Similarly, contemporary hyenas, despite their durophagous nature and formidable bite force, are also known to produce smaller tooth pits (Andrés et al., 2012; Arriaza et al., 2021; Courtenay et al., 2021a). The identification of at least one small tooth pit here has helped our algorithms consider this possible variability, however, the inclusion of larger datasets of undeniable *Pachycrocuta* activity will help refine this aspect of our study. Finally, the poor preservation of cortical surfaces is likely to condition the quality of our predictions, however, considering the abrasive nature of fluvial alterations, it is likely that this will only result in an infra-estimation of the original size of these tooth marks (Gümrükçu and Pante, 2018; Pineda et al., 2023).

The distinctly circular nature of tooth marks here have certainly influenced the morphological affinities that may have been present between *P. brevirostris* and contemporary hyaenids, such as *Crocuta crocuta*, which tend to produce more elongated and superficial pits (Courtenay et al., 2021a, 2023b). Likewise, *P. brevirostris* is thought to be closer phylogenetically to modern species of the genus *Hyaena*, such as the striped (*H. hyaena*) and brown (*H. brunnea*) hyena (Palmqvist et al., 2011). From this perspective, it would be interesting to see how these pits compare with other hyaenid species, recognised for producing smaller and slightly more circular pits than the modern hyaenid species used in the present dataset (Arriaza et al., 2021). It is also important to point out that prior attempts to identify *Pachycrocuta* activity at the palaeontological site of Venta Micena 3 (Yravedra et al., 2022), and archaeological site of Barranco León (Courtenay et al., 2023b), associated much smaller pits than those observed here to *Pachycrocuta*. While these instances might still represent plausible *Pachycrocuta* activity, expanding the reference dataset to encompass unequivocal examples of hyaenid-induced bone damage could offer valuable insights into the precise nature of *Pachycrocuta* bite damage. Likewise, these methods are limited to predicting the most likely carnivoran to have produced the tooth marks based solely on the morphology of tooth pits, with a certain level of uncertainty conditioned by our inability to directly observe these extinct carnivorans. The present study provides additional taphonomic evidence to strongly indicate that the only likely carnivoran to have produced these pits is the *Pachycrocuta*, thus overcoming this limitation, facilitating future predictions of *Pachycrocuta* activity in other sites.

Because the fossil sample is too limited to characterise within-taxon variability empirically, simulations offer a practical way to approximate the expected range of morphological expression and to provide the statistical structure required for quantitative comparison. These simulated variants do not replace the original observations, but they allow the FN3 marks to be contextualised within a broader space of plausible *Pachycrocuta* tooth-mark morphologies. As with any modelling exercise, they represent informed approximations rather than direct observations, but they capture the variation necessary for downstream analyses.

The presentation of the FN3-11-T93-5-1 specimen here, and analysis using artificially intelligent algorithms, therefore provides a new potential means of understanding *Pachycrocuta* tooth marks. This methodological approach has demonstrated its utility in characterising tooth marks made by an extinct carnivore species, a task that would have traditionally been hindered by challenges in taphonomic equifinality in the fossil record. While the results shed light on diagnostic criteria to identify the activity of this carnivoran species, this also highlights the potential computational techniques have to augment palaeontological and archaeological data and research. Such advancements, though promising, necessitate continued refinement and validation to further enhance our insights into the evolutionary history of carnivorans, as well as the interaction and competition they may have had with our own species.

## Acknowledgements

The present study was carried out with the authorisation granted by the Ministry of Education, Culture and Sport of the Junta de Andalucia. Permits were granted in association with the project titled “Primeras ocupaciones humanas y context paleoecológico a partir de los depósi-tos pliopleistocénicos de la Cuenca de Guadiz-Baza. Zona arqueológica de la Cuenca de Orce, Orce, Granada”, coordinated through the University of Granada. Research at Fuente Nueva 3 is currently possible thanks to the support and approval of the Consejería de Turismo, Cultura y Deporte (Junta de Andalucía, Spain) through the General Research Project (2023-2026) “Evolución humana y paleoecología a partir de los yacimientos pleistocenos de la Zona Arqueológica ‘Cuenca de Orce’. Retos y desafíos” (Ref: SIDPH/DI/MCM). The corresponding author wishes to thank the helpful comments and suggestions of Antoine Souron and Jean-Renaud Boisserie, regarding the taxonomic characterisation of hippopotami, Bisrat Araya for their support, as well as three reviewers for their constructive comments. We would like to thank the *Muséum National d’Histoire Naturelle* (MNHN), Paris, for access to the large mammal specimens we inspected during December, 2024, while the corresponding author would also like to thank Lila Geis for her help during this trip. We would like to thank Francesco D’Errico for providing access to the microscopes and equipment used to carry out this study. The corresponding author is also incredibly grateful to the advice and recommendations of Rosa Huguet on all aspects of this research, and Dario Herranz-Rodrigo for his work and advice on multiple occasions. We would also like to thank María Higueras Muñoz for her help in taking photos of the specimens, and the Archaeological and Ethnological Museum of Granada for their help in facilitating access to these specimens. Next we wish to acknowledge the work and efforts of reviewers Sebastián Yrarrázaval, Matheiu Quenu and James Mulqueeney, as well as PCI recommender Anastasia Eleftheriadou for handling our manuscript. Finally, we wish to acknowledge all individuals who have actively participated in the excavations of the Orce sites, both from old campaigns and new ones, as well as the conservation-restauration specialists involved in the handling of such materials.

A Preprint version of this article has been peer-reviewed and recommended by Peer Community In Archaeology (https://doi.org/10.24072/pci.archaeo.100544).

## Fundings

This work was supported by the Consejería de Turismo, Cultura y Deporte [grant reference SIDPH/DI/MCM], and the project PID 2021.125098NB.I00 by the MCIN / AEI / 10.13039 / 501100011033 / FEDER Una Manera de Hacer Europa. This research was also supported by the Spanish Ministry of Science and Innovation through the “María de Maeztu” excellence accreditation (CEX2024-01485-M/funded by MICIU/AEI/10.13039/501100011033) and the project Cognitive processes of the first humans: dialectic between technology and paleoenvironment in the circum-Mediterranean area from the Late Pliocene to the Middle Pleistocene (HUMANCOGNENV) (PID2024-156295NB-I00). We also recieved funding from the Generalitat de Catalunya Research Groups (AGAUR) IPHES-CERCA and URV Paleoecology of Pliocene and Pleistocene and Human Dispersals (PalHum) 2021 SGR 01238. J.S. was funded by the Research Council of Finland (project nr. 340775) during this work. L.A.C. was originally funded by the ERC Synergy project QUANTA (grant number 951388) and the French government in the framework of the University of Bordeaux’s IdEx “Investments for the Future” program / GPR “Human Past”. L.A.C. is currently funded by the Agence Nationale de la Recherche, with an Access-ERC project titled BSMART (ANR-25-AERC-0005).

## Author Contributions

L.A.C. Conceptualization, Methodology, Software, Validation, Formal Analysis, Investigation, Resources, Data Curation, Writing - original draft, review and editing, Visualization. A.S.R. Resources. J.S. Validation. S.V. Validation. D.B. Supervision. J.M.J.A. Resources, supervision, project administration, funding acquisition. J.Y. Investigation, supervision, project administration.

## Conflict of interest disclosure

The corresponding author would like to state that they have no competing interests to declare.

## Data, script, code, and supplementary information availability

### Data Availability Statement

Some data has also been included in the following repository; https://github.com/LACourtenay/VAE_MCMC_Pachycrocuta_Simulations, however the majority of the reference data has not been included as they come from other studies, that are also open access, and have been suitably cited within the main text of the present study. In the case of (Courtenay et al., 2021b), the present authors have acquired the necessary permissions to save the dataset to the repository where our code is included. Nevertheless, the code presented in the repository associated with the present paper, however, includes some lines that are used to load into memory the data from these studies, so that they can then be used in subsequent analyses.

### Code Availability Statement

All code used for the present study are available on the corresponding author’s GitHub page at; https://github.com/LACourtenay/VAE_MCMC_Pachycrocuta_Simulations.

### Supplemental Information

Supplementary File 1 has been included at https://github.com/LACourtenay/VAE_MCMC_Pachycrocuta_Simulations. This file contains - Supplementary Text: Site Presentation and Imbalance Calculations; Supplementary Table 1; Supplementary Figure 1 to 12

## Notes

https://github.com/LACourtenay/VAE_MCMC_Pachycrocuta_Simulations

### Competing Interest Statement

The authors have declared no competing interest.

### Summary of Updates

This updated submission is to now include the URL and appropriate badge linking the paper with its PCI Archaeology recommendation that can be found at https://doi.org/10.24072/pci.archaeo.100544

