## Supplementary Figures and Appendices for "Computational simulations of potential *Pachycrocuta* bite damage based on a ∼1.2 Ma ravaged hippopotamus femur from Fuente Nueva 3 (Orce, Granada, Spain)"

Supplmenetary Materials

**Lloyd A. Courtenay**
Préhistoire à l'Actuel Culture, Environment et Anthropologie (PACEA UMR 5199)
University of Bordeaux & CNRS, France
Department d'Historia i Historia de l'Art, Universitat Rovira i Virgili (URV), Tarragona, Spain


**Alexia Serrano-Ramos**
Institut Català de Paleoecologia Humana I Evolució Social (IPHES-CERCA)
& Department of Prehistory and Archaeology, University of Granada, Spain

**Juha Saarinen**
Department of Geosciences and Geography, University of Helsinki, Helsinki, Finland

**Suvi Viranta**
Department of Anatomy, University of Helsinki, Helsinki, Finland

**Deborah Barsky**
Institut Català de Paleoecologia Humana I Evolució Social (IPHES-CERCA)
& Department d'Historia i Historia de l'Art, Universitat Rovira i Virgili (URV), Tarragona, Spain

**Juan Manuel Jiménez-Arenas**
Department of Prehistory and Archaeology
& Excellence Unit Archaeometrical Studies, University of Granada, Spain

**José Yravedra**
Department of Prehistory, Ancient History and Archaeology
& C.A.I. Archaeometry and Archaeological Analysis, Complutense University, Madrid, Spain

December 16, 2025

#### Abstract

The following document contains all of the supplementary information related to our paper "Computational simulations of *Pachycrocuta* bite damage based on a ∼1.2 Ma ravaged hippopotamus femur from Fuente Nueva 3 (Orce, Granada, Spain)". This includes supplementary figures, an extended version of Table 1 from the main paper, as well as supplementary text with site presentation details.

***Keywords*** Taphonomy · Geometric Morphometrics · Unsupervised Learning · Variational Autoencoders · Markov Chain Monte Carlo · Confocal Microscopy · Numeric Simulation

### Contents

**1 Appendix A: Site Presentation of Fuente Nueva 3 (FN3, Orce, Granada, Spain) 3**

**2 Appendix B: Assessing the Impact of Imbalance on VAE Training 4**

**3 Appendix C: Supplementary Tables and Figures 6**
3.1 Supplementary Figures . . . . . . . . . . 6
3.2 Supplementary Tables . . . . . . . . . . 15

**References 17**

### 1 Appendix A: Site Presentation of Fuente Nueva 3 (FN3, Orce, Granada, Spain)

The site of Fuente Nueva 3 (FN3) is located in the north-east of the Guadix-Baza Basin (Granada, Spain). FN3 is located in proximity to a number of major archaeological and palaeontological localities, including the sites of Venta Micena 3 and 4 (Martínez-Navarro and Palmqvist, 1995; Luzón et al., 2021), as well as the site of Barranco León, the latter of which has yielded a decidual molar attributed to *Homo* sp., one of the oldest hominin fossils of western Europe (Oms et al., 2000; Toro-Moyano et al., 2013). Discovered in 1991, FN3 is an open-air site with an excavated surface area of 104 $m^2$, situated in sedimentary deposits on the edge of the Baza palaeolake. FN3 is situated on the Upper Member of the Baza Formation, characterised by distal fluvial/alluvial sedimentation (Calvache and Viseras, 1997). Evidence of invertebrate fauna reveal an association of this site with alternating shallow aquatic and areal environments (Anadón et al., 2003).

A total of 8 stratigraphic geological layers have been described in FN3, 3 of which contain archaeo-paleontological deposits of interest, denoted as layers 2, 3 and 5 (Oms et al., 2000, 2010, 2011). FN3-5 presents the most abundant accumulation of archaeological artefacts, dated at $\approx$1.2 Ma via a combination of magnetostratigraphic and biochronological data, alongside U-series and ESR dating methods (Duval et al., 2012). FN3-5 consists of fine-grained greenish sands and mudstones, which is further subdivided into subunits 5A and 5B. No taphonomic data reveals the reworking or resedimentation of these units, while many skeletal elements of large herbivores from this layer have been noted to have been preserved in anatomical connection.

FN3-5 presents a rich fossil assemblage of large herbivores. A complete list of these species is presented as follows; *Mammuthus meridionalis*, *Stephanorhinus* cf. *etruscus*, *Equus altidens*, *Equus sussenborensis*, *Hippopotamus antiquus*, *Bison* sp. *Hemitragus* cf. *albus*, *Praemegaceros* cf. *verticornis*, and *Metacervocerus rhenanus* (Martínez-Navarro et al., 2010; Yravedra et al., 2021; Alberdi, 2010; Abbazzi, 2010). FN3 is additionally characterised by a large number of carnivoran species; *Ursus etruscus*, *Canis mosbachensis*, *Xenocyon lycaonoides*, *Vulpes alopecoides*, *Meles meles*, *Martellictis ardea*, *Pachycrocua brevirostris*, *Lynx* cf. *pardina* and an indeterminable species of Felidae (Martínez-Navarro et al., 2010; Medin et al., 2017; Bartolini-Lucenti and Madurell-Malapiera, 2020,?; Madurell-Malapeira et al., 2011; Ros-Montoya et al., 2021; Boscaini et al., 2015). Although not directly detected in the FN3 assemblage, the Orce paleolandscape has also revealed the presence of other carnivoran species, including; *Megantereon cultridens/whitei*, *Panthera gombaszoegensis*, *Homotherium* sp. and *Acinonyx pardinensis* (Turner, 1995, 1992; Turner and Antón, 1997; Antón et al., 2005; Antón, 2013; Martínez-Navarro and Palmqvist, 1995).

The micromammal assemblage of FN3-5 consists of the following; *Mimomys savini*, *Manchenomys orcensis*, *Allophaiomys* aff. *lavocati*, *Apodemus* aff. *sylvaticus*, *Castillomys rivas*, *Discoglossus* cf. *jeanneae*, *Pelobates cultripes*, *Bufo* gr. *B. bufo*, *Pelophylax* cf. *perezi*, *Malpolon monspessulanus*, *Natrix maura*, Lacertidae indet, cf. *Coronella* sp. and *Natrix* indet (Agustí et al., 2010, 2022; Blain, 2005, 2009; Blain and Bailon, 2010; Blain et al., 2011, 2016; Furió, 2007).

Ecomorphological data derived from both micromammal and large mammal data reveal FN3-5 to have been deposited in a relatively dry and cold climate, with a mean annual temperature of 14.7 °, and mean annual precipatation of 618 mm (Saarinen et al., 2021; Sánchez-Bandera et al., 2023; Agustí et al., 2022). The vegetation of this landscape would have most likely consisted in open vegetation with pines, junipers, oaks, and wild olive, alongside a diversity of woody taxa (Ochando et al., 2022).

Taphonomically, the FN3-5 assemblage is characterised by a high degree of fragmentation, with generally low degrees of weathering, alterations produced by water currents, and biochemical processes. The impact of both carnivorans and humans have been generally characterised as being low (Yravedra et al., 2021; Espigares et al., 2019).

The stone tool assemblage from FN3 was knapped from local raw materials and is characterized by numerous small flint flakes, some of which were knapped using the bipolar-on-anvil method, as well as abundant limestone percussion tools, typical of the Oldowan techno-complex in Europe (Toro-Moyano et al., 2010, 2011; Barsky et al., 2010, 2013, 2015a,b, 2018, 2022).

#### 2 Appendix B: Assessing the Impact of Imbalance on VAE Training

Following the recommendations and inquiries of the reviewers, this section assesses the potential impact of imbalanced datasets on our results.

The datasets used in the present study vary substantially in sample size, with the smallest sample belonging to *Vulpes vulpes* (n = 53) and the largest to *Canis lupus* (n = 283). This disparity yields a balance index of 0.92 when considering all labels, but in the extreme case of comparing *C. lupus* with *V. vulpes*, the index drops to 0.68. The balance index is computed as:

$$B = \frac{-\sum_{i=1}^{g} \frac{c_i}{n}; \log \frac{c_i}{n}}{\log; g}, \qquad B \in [0, 1] \tag{1}$$

adapted from Shannon (1948a,b), where $c_i$ is the count of individuals in group $i$, $g$ is the total number of groups, and the sum of all $c_i$ equals $n$.

Class imbalance occurs when the different classes in a labeled dataset are represented by highly unequal sample sizes. Such imbalance can bias learning algorithms because the majority class dominates the training process, potentially leading to poor generalization on minority classes.

In supervised learning, models aim to minimize classification error based on class labels. When one class is overrepresented, algorithms often fail to capture the variability of minority classes, producing noisier estimates and biased predictions.

Unsupervised learning, by contrast, does not use class labels to update model parameters. Its objective is to approximate the joint distribution of the data across all samples. From this perspective, unsupervised learning is less sensitive to class imbalance, except in cases where groups occupy distinct, non-overlapping regions of feature space. In such cases, the underlying distribution may still be influenced by sample sizes.

In this study, we aim to model the overall distribution of carnivore traits rather than individual species. Consequently, the potential effect of sample imbalance was initially considered negligible. Nevertheless, as suggested by reviewers, overrepresented species naturally contribute more to the latent distributions, which could influence the learning process and the resulting simulations. Therefore, a critical assessment is whether overlap in latent space is independent of raw species counts. If distributions are largely overlapping, then imbalance does not distort the general variability captured by the algorithm, and the simulations can be considered reliable.

The VAE's loss function minimizes the KL divergence between each species' latent distribution and a prior Gaussian distribution. If the latent dimensions do not fulfill this objective, it would indicate that the model has failed to capture the intended distribution, rendering the results unreliable.

To investigate this, we examine whether there are substantial differences in KL divergence between samples. The null hypothesis is that if latent space is conditioned by label distributions, then samples will not have converged to the prior distribution. Since all species have sample sizes sufficient to satisfy the central limit theorem, the covariance matrices of each latent distribution can be reliably estimated.
The pairwise KL divergence between two samples is defined as:

$$D_{KL}\left(\mathcal{N}_1 | \mathcal{N}_2\right) = \frac{1}{2}\left[\text{tr}\left(\Sigma_2^{-1}\Sigma_1\right) - k + \ln\frac{\det\Sigma_2}{\det\Sigma_1}\right] \tag{2}$$

$$D_{KLsim} = \frac{1}{2} \times D_{KL}\left(\mathcal{N}_1 | \mathcal{N}2\right) + D_{KL}\left(\mathcal{N}_2 | \mathcal{N}_1\right) \tag{3}$$

where $\mathcal{N}$ is the normal distribution modeled by the VAE, $\Sigma$ is the covariance matrix of each group, and $k$ is the dimensionality (here, 15). For this study, we calculated the KL divergence between all pairs of species to identify any substantial differences.

To determine what constitutes an acceptable divergence, we performed simulations of distribution pairs in the same latent space, gradually increasing their differences. This produced a calibration curve indicating how the KL metric behaves as distributions become more distinct.

The results of these simulations and their comparison with the real data are summarized in Figure 6 of the main text. As the mean shift increases from 0 (identical distributions) to 10 (drastically different distributions), the KL metric grows roughly quadratically with the Mahalanobis distance. The KL values obtained from our carnivore samples fall approximately on the curve corresponding to a mean shift of about 1, with most values ranging between 30 and 60. The curve itself only begins to show substantial divergence at KL $\approx$ 200, indicating that the observed species distributions represent only moderate differences.

In a 15-dimensional Gaussian latent space, a mean-shift of 1 corresponds to differences of roughly one standard deviation along each dimension. This demonstrates that the variability captured by the VAE is relatively minor and reflects general variation rather than structural differences between species. The exception is wolves and lions, which present the highest divergence (KL = 311.4), representing more pronounced differences in latent space.

To assess whether sample imbalance might influence these results, we examined the correlation between species KL values and sample size. The Spearman correlation was low ($\rho = 0.21$, $p = 0.27$, FPR = $49\%$), suggesting that sample size does not notably bias the latent distributions. From this perspective, class imbalance is unlikely to have affected the overall results of the present study.

### 3 Appendix C: Supplementary Tables and Figures

#### 3.1 Supplementary Figures

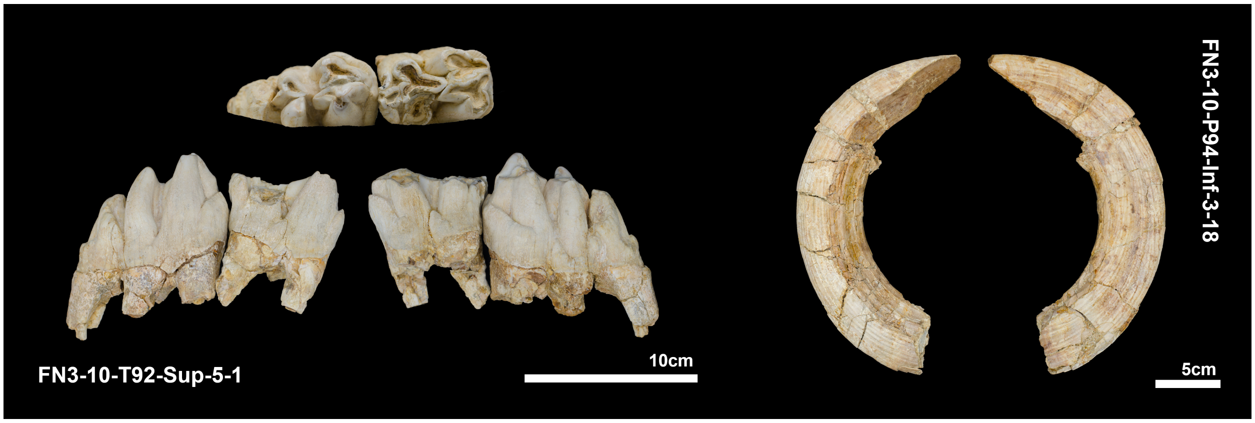


**Supplementary Figure 1:** Examples of dental remains of *H. antiquus* recovered from the site of Fuente Nueva 3. Photos were taken by María Higueras Muñoz

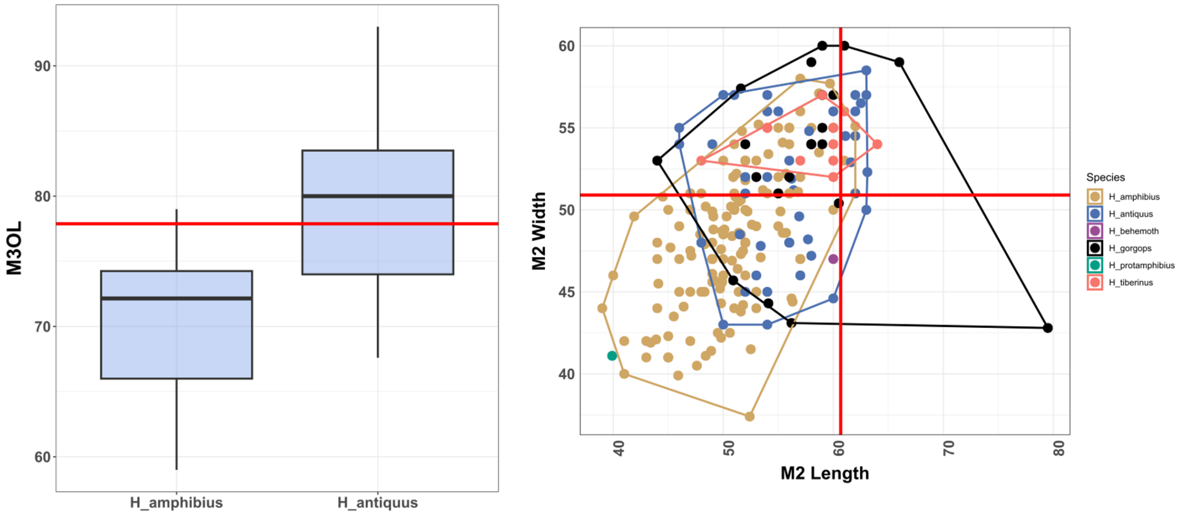


**Supplementary Figure 2:** Results comparing the metric properties of Hippopotamus teeth (FN3-10-T92-Sup-5-1) recovered from Level 5 of FN3 with multiple comparative samples. $M_3$OL refers to the occlusal length of the lower 3rd molar, compared with data published by Mecozzi et al. Mecozzi et al. (2023). Length and widths of the lower second molar were compared with published data by Fidalgo et al. Fidalgo et al. (2023).

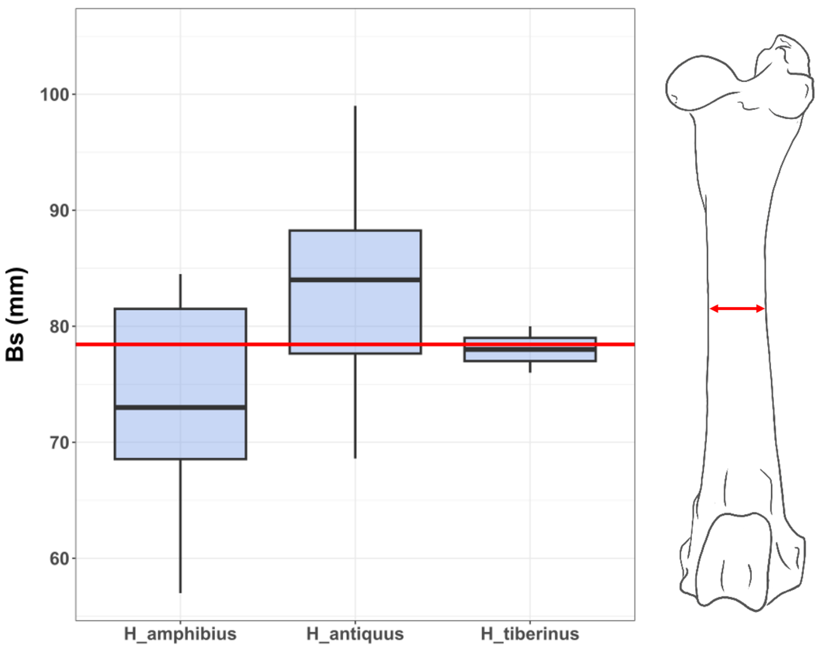


**Supplementary Figure 3:** Results comparing the width (Bs) of the FN3-11-T93-Sup-5-1 diaphysis with comparative samples and measurements of multiple species of hippopotamus provided by Fidalgo et al. Fidalgo et al. (2023).

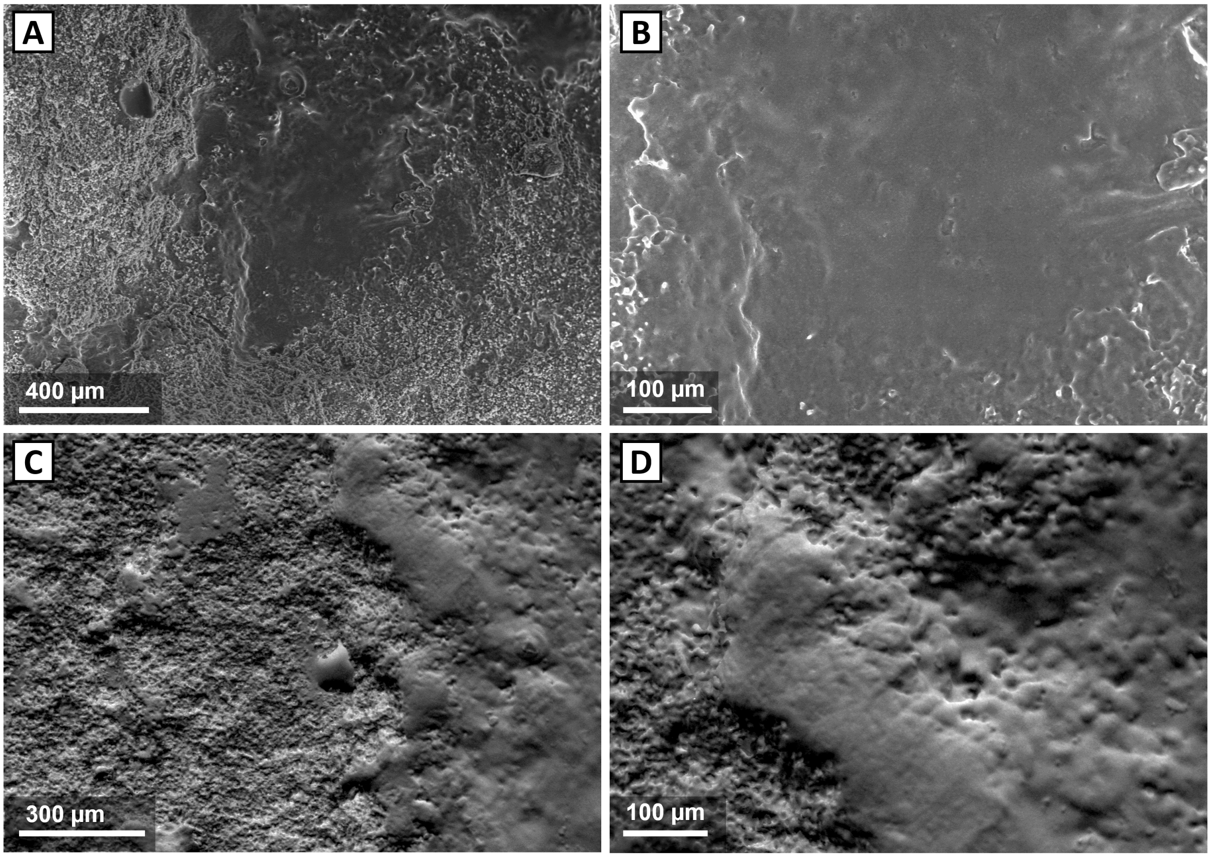


**Supplementary Figure 4:** Scanning Electron Microscope (SEM) documentation of multiple areas of taphonomic interest of the FN3-11-T93-Sup-5-1 diaphysis. Note the polished nature of certain areas of the cortical surface due to fluvial alterations, while a number of other areas occupying the majority of the cortical surface present extreme levels of abrasion. Panels (B) and (D) are details of panels (A) and (C).

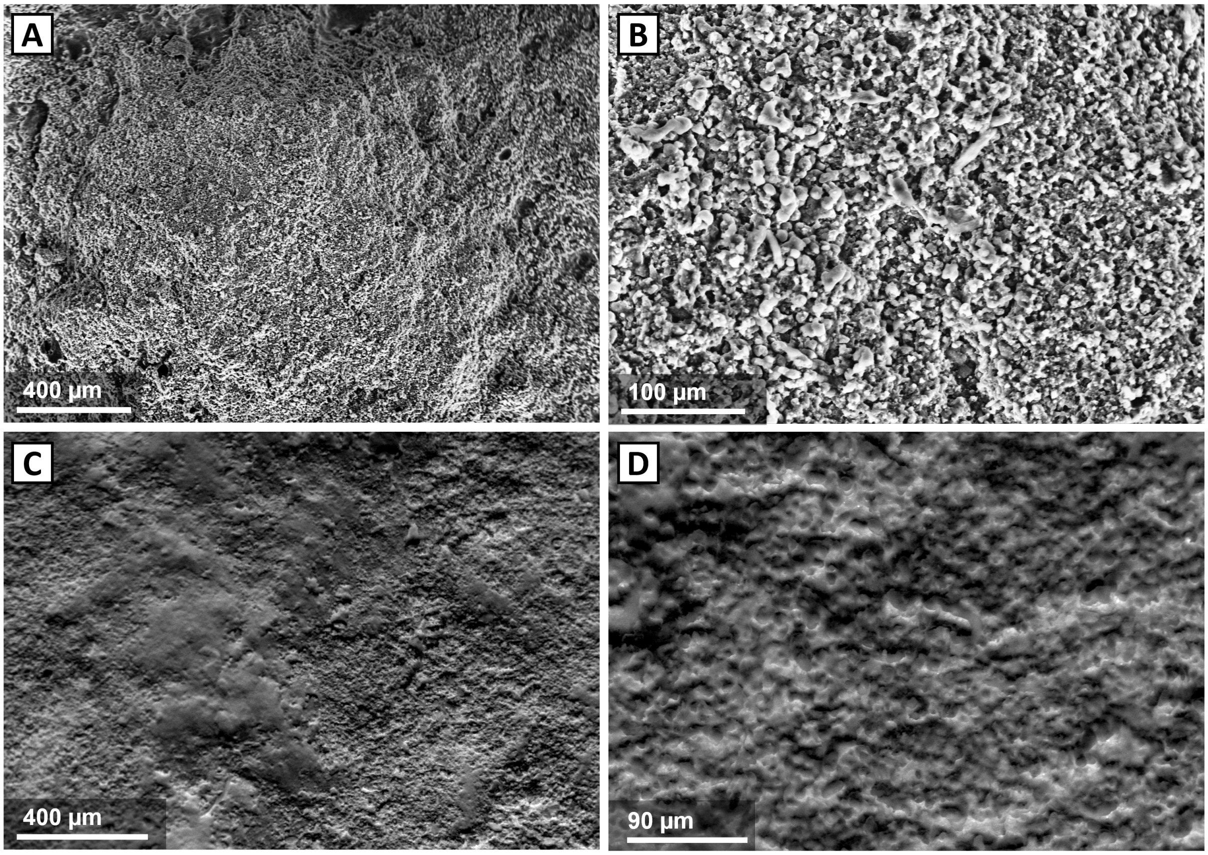


**Supplementary Figure 5:** Scanning Electron Microscope (SEM) documentation of multiple areas of taphonomic interest of the FN3-11-T93-Sup-5-1 diaphysis. These images detail the more extreme instances where fluvial activity have completely abraded and obliterated the cortical surface. Panels (B) and (D) are details of panels (A) and (C).

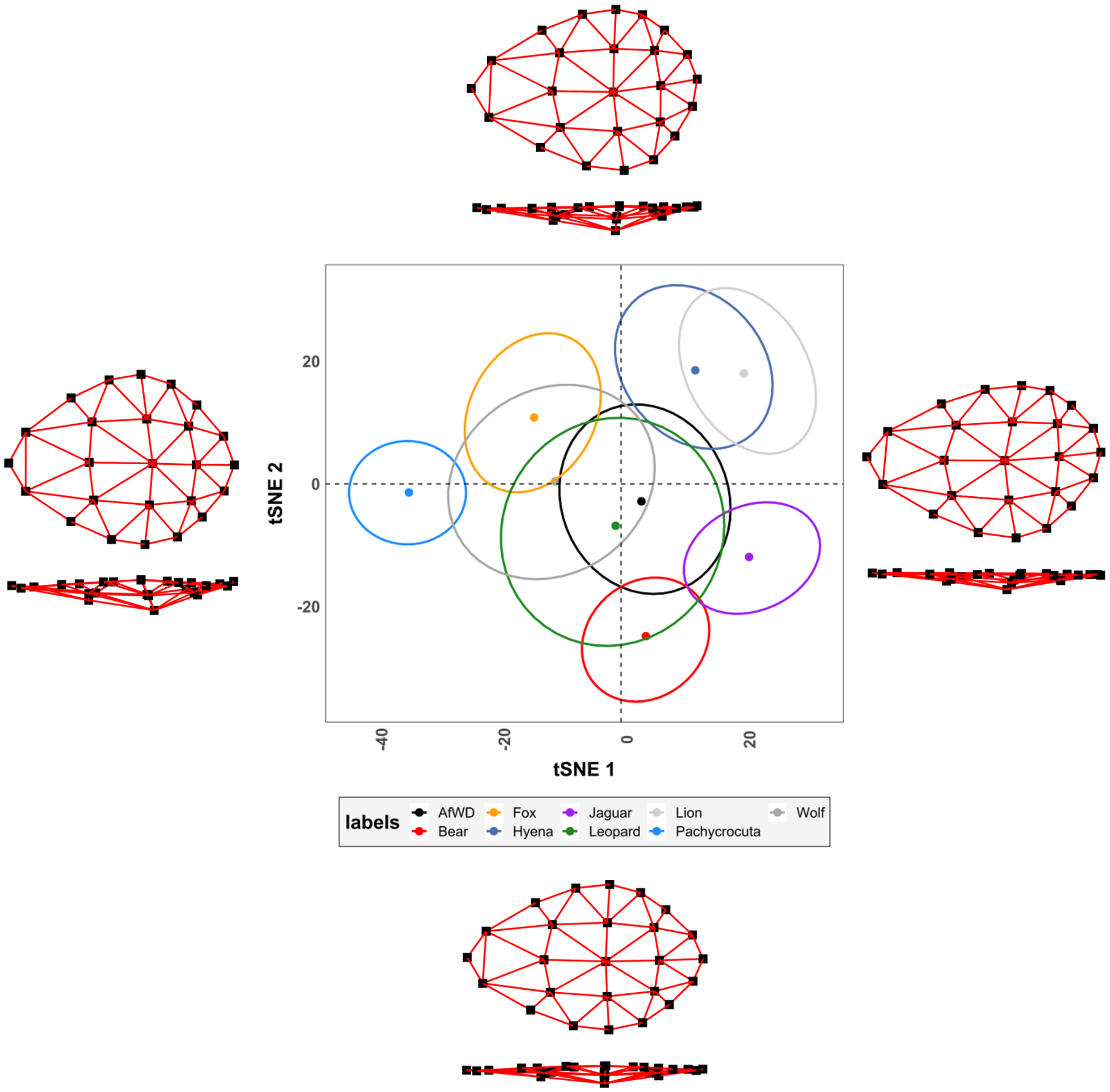


**Supplementary Figure 6:** t-Distributed Stochastic Neighbour Embeddings (tSNE) of the remaining morphological shape variability simulated by Variational Autoencoders (VAE) for the median specimen of each species. Morphological variability across feature space is represented on the extremity of each tSNE score

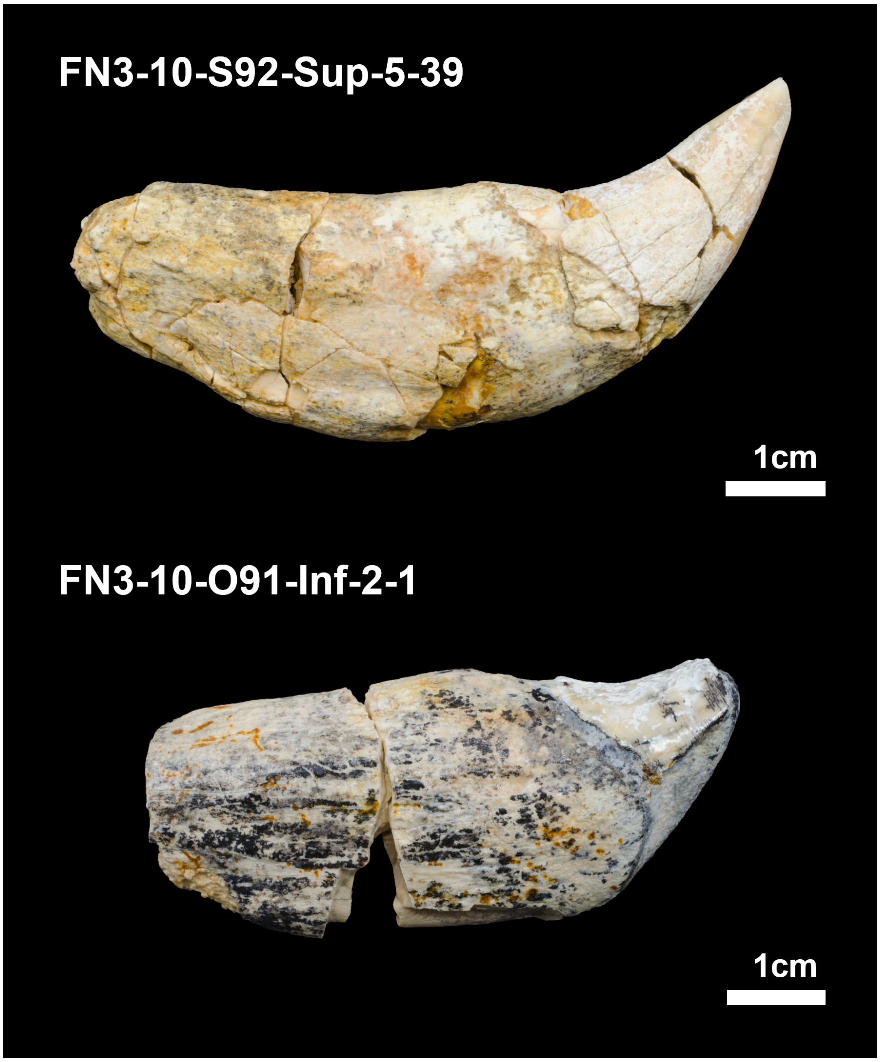


**Supplementary Figure 7:** Examples of dental remains of *Pachycrocuta brevirostris* recovered from the site of Fuente Nueva 3. Photos were taken by María Higueras Muñoz

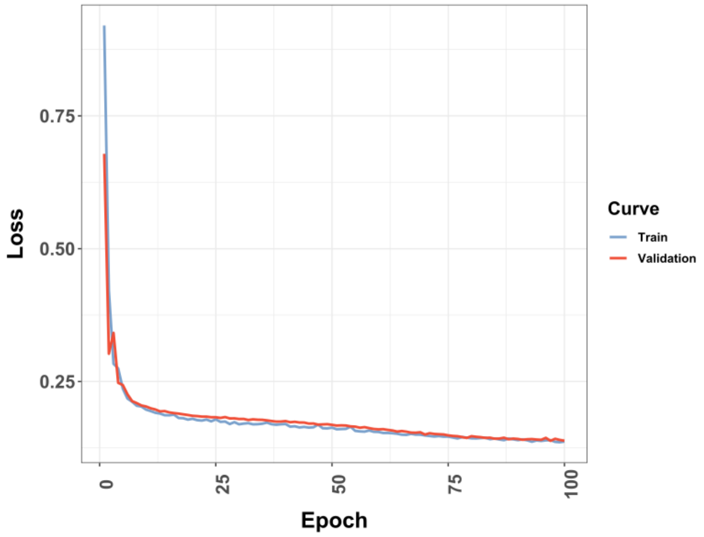


**Supplementary Figure 8:** Variational Autoencoder (VAE) learning curves for training and validation data.

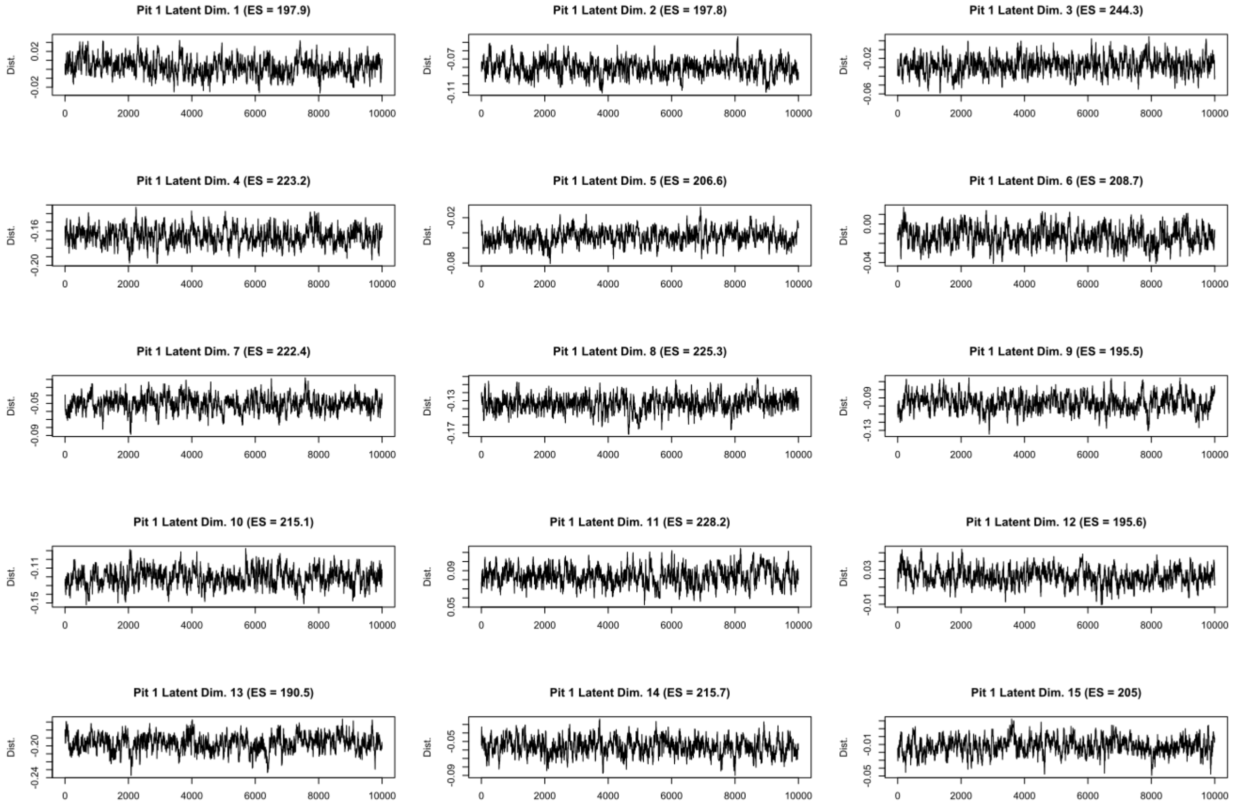


**Supplementary Figure 9:** Markov Chain Monte Carlo trace evaluation for each of the latent dimensions when sampling based on the first pit identified on the FN3-11-T93-Sup-5-1 individual. ES indicates the Effect Size metric used to evaluate the trace, with values above 100 being considered optimal.

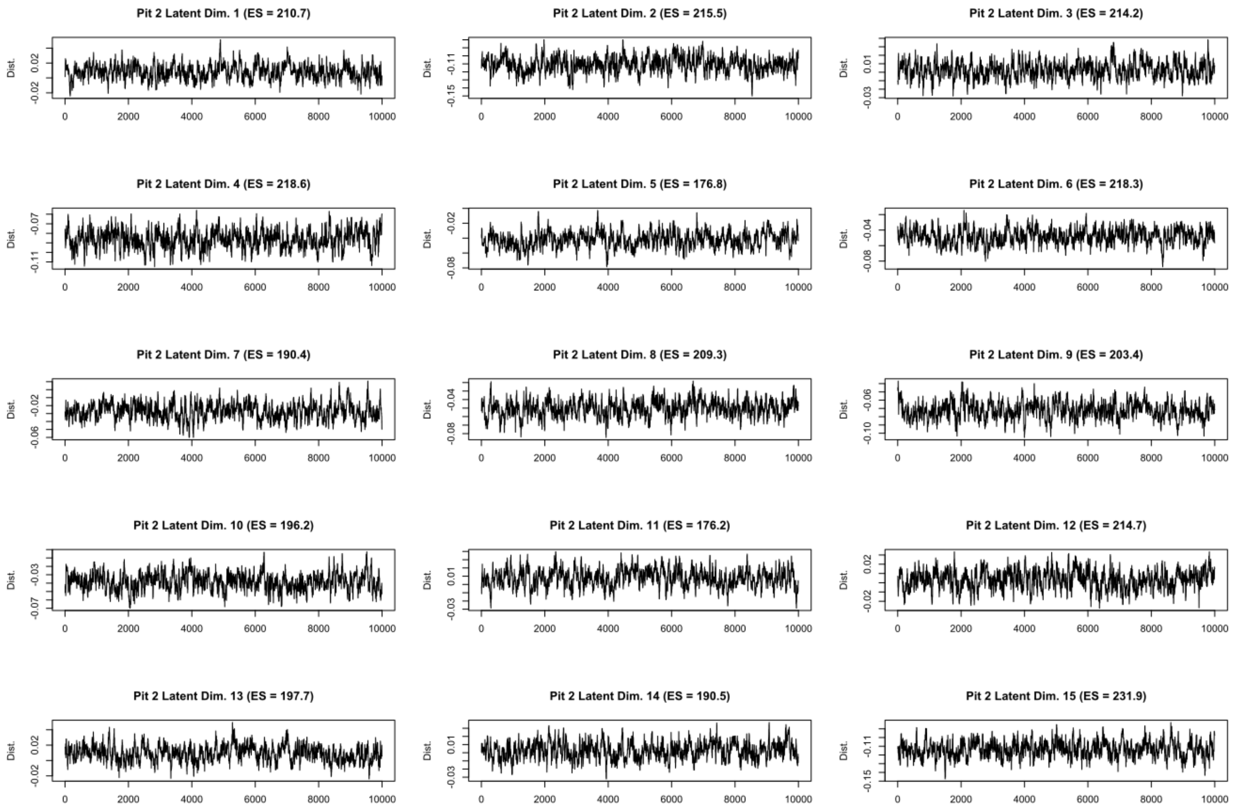


**Supplementary Figure 10:** Markov Chain Monte Carlo trace evaluation for each of the latent dimensions when sampling based on the second pit identified on the FN3-11-T93-Sup-5-1 individual. ES indicates the Effect Size metric used to evaluate the trace, with values above 100 being considered optimal.

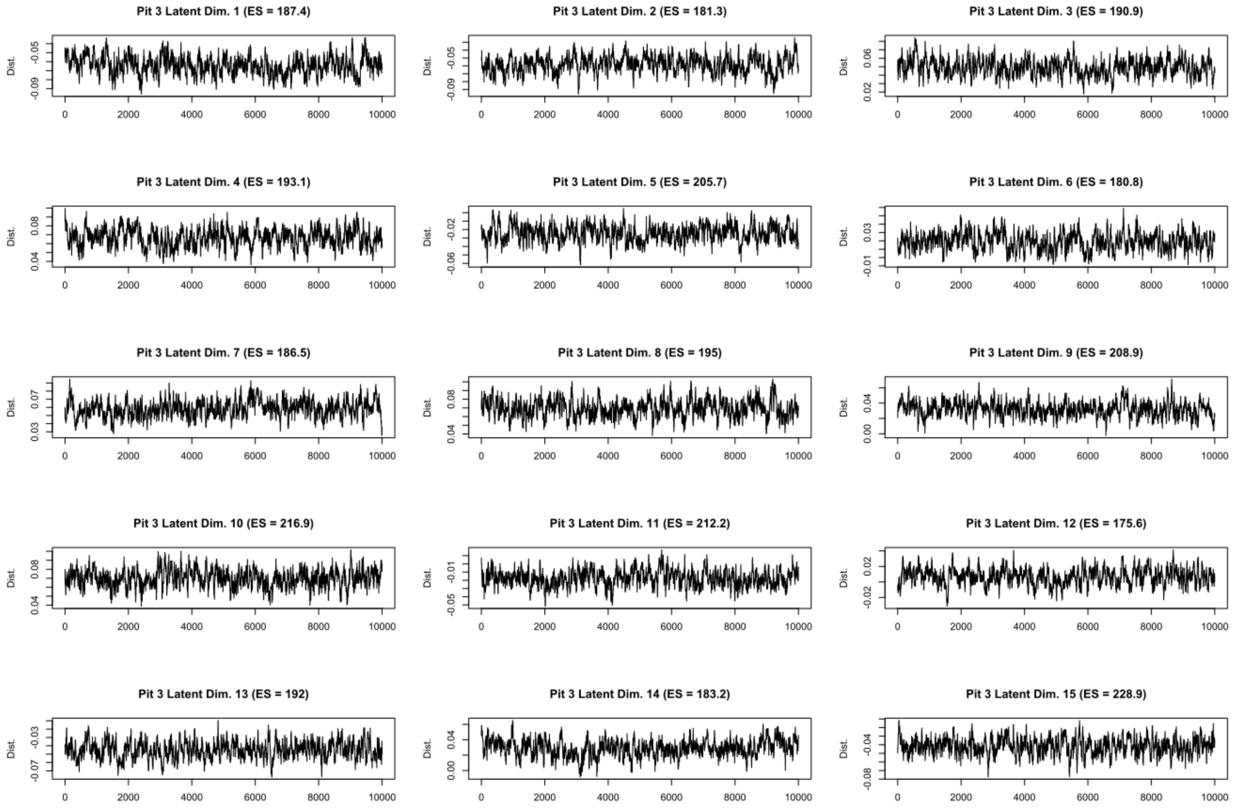


**Supplementary Figure 11:** Markov Chain Monte Carlo trace evaluation for each of the latent dimensions when sampling based on the third pit identified on the FN3-11-T93-Sup-5-1 individual. ES indicates the Effect Size metric used to evaluate the trace, with values above 100 being considered optimal.

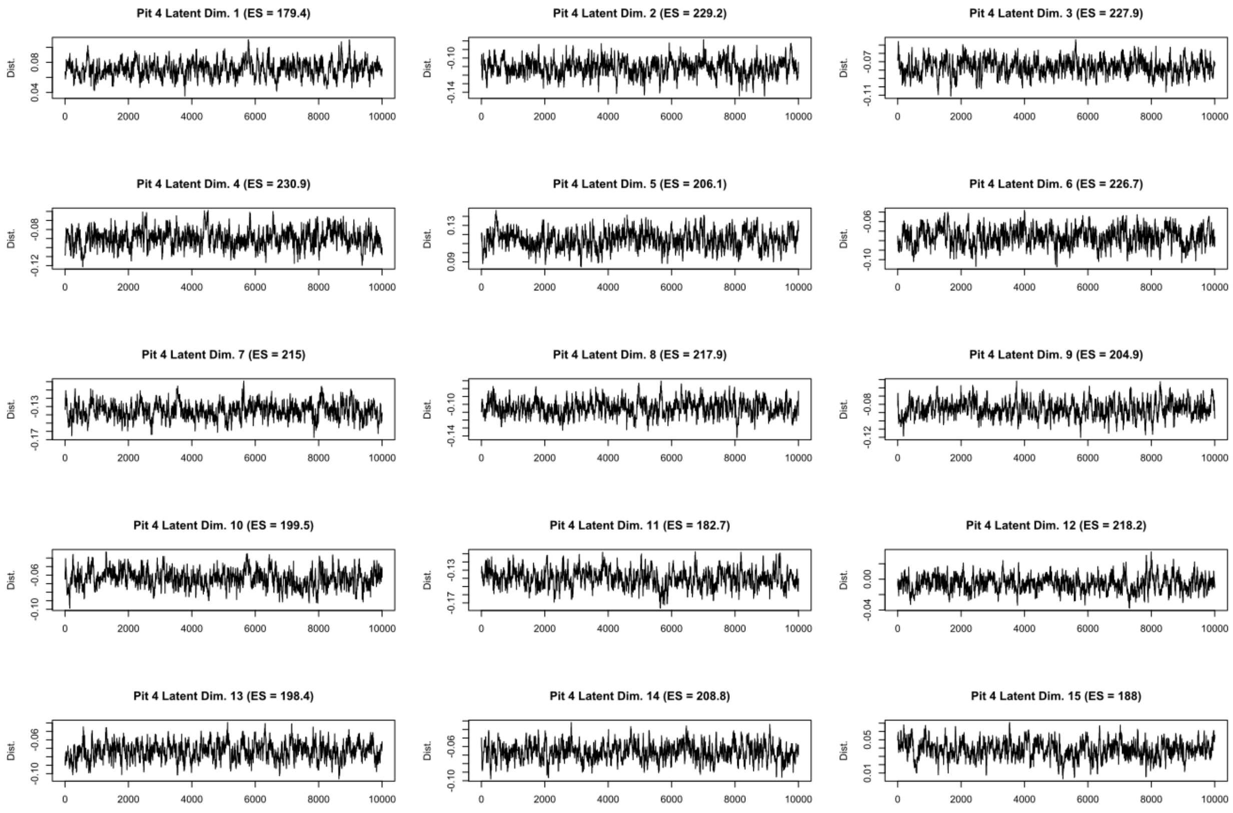


**Supplementary Figure 12:** Markov Chain Monte Carlo trace evaluation for each of the latent dimensions when sampling based on the fourth pit identified on the FN3-11-T93-Sup-5-1 individual. ES indicates the Effect Size metric used to evaluate the trace, with values above 100 being considered optimal.

##### 3.2 Supplementary Tables

The following reference list is relevant for the content of Supplementary Table 1; Shapiro and Wilk (1965); Royston (1995); Höhle and Höhle (2009); Hasan et al. (2011); Razali (2011); Rodríguez-Gonzálvez et al. (2014); Nocerino et al. (2017); Herrero-Huerta et al. (2018); Ariza-López et al. (2019); Benjamin and Berger (2019); Colquhoun (2019); Rodríguez-Martín et al. (2019a,b); Courtenay et al. (2020, 2021)

**Supplementary Table 1:** Descriptive statistics of the measurements extracted from tooth pit samples, including the simulated measurements of possible marks by *Pachycrocuta brevirostris*. Measurments are all reported in mm. Central Tendency values are reported as either the mean or the median, depending on whether the sample distribution were found to be homogeneous or not, with the mean being used for homogeneous distributions, and the median used for inhomogeneous distributions. Likewise, deviation values were either reported as the standard deviation or the Square Root of the Biweight Midvariance for the same reasons . Determining whether distributions were homogeneous or not were carried out using the Shapiro-Wilks test, with the results for said test reported including the test statistic W, the $p$ values, as well as the probability of this $p$-value leading to a False Positive (False Positive Risk; FPR). FPR values are reported in percentages, and were calculated using 0.5 as a prior. Robust statistical metrics were therefore used for values of $p < 0.003$. Upper and Lower bounds of Confidence Intervals (CI) were calculated using 95% bootstrapped intervals.

| | | W | $p$ | FPR (%) | Min. | Lower CI | Central | Deviation | Upper CI | Max. |
|---|---|---|---|---|---|---|---|---|---|---|
| *V. vulpes* | Length | 0.95 | 0.020 | 17.86 | 0.520 | 1.730 | 1.976 | 0.892 | 2.222 | 3.941 |
| | Width | 0.94 | 0.01 | 11.08 | 0.501 | 1.12 | 1.41 | 0.593 | 1.520 | 3.209 |
| | Depth | 0.78 | $1.8 \times 10^{-7}$ | $7.7 \times 10^{-4}$ | 0.045 | 0.215 | 0.255 | 0.160 | 0.342 | 1.117 |
| *P. pardus* | Length | 0.88 | $1.3 \times 10^{-6}$ | 0.005 | 1.272 | 1.977 | 2.410 | 0.972 | 2.744 | 6.346 |
| | Width | 0.91 | $3.0 \times 10^{-5}$ | 0.109 | 0.755 | 1.332 | 1.556 | 0.713 | 1.808 | 3.735 |
| | Depth | 0.82 | $7.5 \times 10^{-9}$ | $3.8 \times 10^{-5}$ | 0.038 | 0.222 | 0.266 | 0.207 | 0.347 | 1.493 |
| *C. lupus* | Length | 0.87 | $4.4 \times 10^{-7}$ | 0.002 | 1.09 | 2.227 | 2.410 | 0.986 | 2.591 | 8.294 |
| | Width | 0.91 | $3.0 \times 10^{-5}$ | 0.087 | 0.732 | 1.635 | 1.739 | 0.727 | 1.995 | 4.639 |
| | Depth | 0.67 | $4.0 \times 10^{-12}$ | $2.9 \times 10^{-8}$ | 0.045 | 0.152 | 0.185 | 0.108 | 0.209 | 1.435 |
| *U. arctos* | Length | 0.93 | $7.9 \times 10^{-4}$ | 1.509 | 0.877 | 2.502 | 3.035 | 1.890 | 3.750 | 9.493 |
| | Width | 0.93 | $5.5 \times 10^{-4}$ | 1.117 | 0.496 | 1.584 | 1.975 | 1.172 | 2.562 | 6.695 |
| | Depth | 0.77 | $5.5 \times 10^{-9}$ | $2.8 \times 10^{-5}$ | 0.023 | 0.191 | 0.272 | 0.307 | 0.430 | 2.422 |
| *L. pictus* | Length | 0.87 | $2.5 \times 10^{-4}$ | 0.564 | 1.125 | 3.231 | 3.482 | 1.703 | 4.480 | 12.596 |
| | Width | 0.88 | $8.1 \times 10^{-7}$ | 0.003 | 0.886 | 2.037 | 2.548 | 1.378 | 2.869 | 7.776 |
| | Depth | 0.74 | $3.1 \times 10^{-11}$ | $2.0 \times 10^{-7}$ | 0.059 | 0.384 | 0.497 | 0.442 | 0.612 | 4.160 |
| *C. crocuta* | Length | 0.87 | $5.6 \times 10^{-7}$ | 0.002 | 1.278 | 3.513 | 3.893 | 1.703 | 4.480 | 12.596 |
| | Width | 0.89 | $2.0 \times 10^{-6}$ | 0.007 | 1.096 | 2.495 | 2.776 | 1.290 | 3.097 | 9.444 |
| | Depth | 0.71 | $8.0 \times 10^{-12}$ | $5.5 \times 10^{-8}$ | 0.082 | 0.351 | 0.421 | 0.266 | 0.491 | 3.176 |
| *P. onca* | Length | 0.87 | $1.5 \times 10^{-6}$ | 0.005 | 0.855 | 3.299 | 3.863 | 2.537 | 4.262 | 16.863 |
| | Width | 0.88 | $2.8 \times 10^{-6}$ | 0.009 | 0.621 | 2.093 | 2.437 | 1.713 | 3.092 | 9.947 |
| | Depth | 0.81 | $1.4 \times 10^{-8}$ | $7.0 \times 10^{-5}$ | 0.047 | 0.300 | 0.355 | 0.410 | 0.555 | 2.166 |
| *P. leo* | Length | 0.92 | $1.3 \times 10^{-4}$ | 0.309 | 2.112 | 4.946 | 5.916 | 3.363 | 6.768 | 18.820 |
| | Width | 0.95 | 0.005 | 6.263 | 1.241 | 3.688 | 3.994 | 2.209 | 4.818 | 9.680 |
| | Depth | 0.91 | $2.2 \times 10^{-5}$ | 0.063 | 0.142 | 0.721 | 0.975 | 0.839 | 1.162 | 3.882 |
| *P. brevirostris* | Length | 0.77 | $2.2 \times 10^{-16}$ | $2.2 \times 10^{-12}$ | 1.627 | 5.817 | 6.612 | 2.412 | 7.365 | 8.129 |
| | Width | 0.84 | $2.2 \times 10^{-16}$ | $2.2 \times 10^{-12}$ | 1.589 | 4.546 | 4.556 | 1.613 | 4.566 | 6.533 |
| | Depth | 0.83 | $2.2 \times 10^{-16}$ | $2.2 \times 10^{-12}$ | 0.236 | 0.886 | 0.910 | 0.274 | 0.925 | 1.162 |
